# Profound phenotypic and epigenetic heterogeneity of the HIV-1 infected CD4+ T cell reservoir

**DOI:** 10.1101/2022.04.02.486753

**Authors:** Vincent H. Wu, Son Nguyen, Jayme M.L. Nordin, Jaimy Joy, Felicity Mampe, Pablo Tebas, Luis J. Montaner, Katharine J. Bar, Laura A. Vella, Michael R. Betts

**Affiliations:** Department of Microbiology, Perelman School of Medicine, University of Pennsylvania (Philadelphia, PA, USA); Department of Medicine, Perelman School of Medicine, University of Pennsylvania (Philadelphia, PA, USA); Center for AIDS Research, University of Pennsylvania (Philadelphia, PA, USA); The Wistar Institute (Philadelphia, PA, USA); Division of Infectious Diseases, Children’s Hospital of Philadelphia (Philadelphia, PA, USA)

## Abstract

Understanding the complexity of the long-lived HIV reservoir during antiretroviral therapy (ART) remains a major impediment for HIV cure research. To address this, we developed single-cell viral ASAPseq to precisely define the unperturbed peripheral blood HIV-infected memory CD4+ T cell reservoir from antiretroviral treated people living with HIV (ART-PLWH) via the presence of integrated accessible proviral DNA in concert with epigenetic and cell surface protein profiling. We identified profound reservoir heterogeneity within and between ART-PLWH, characterized by novel and known surface markers within total and individual memory CD4+ T cell subsets. We further uncovered novel epigenetic profiles and transcription factor motifs enriched in HIV-infected cells that suggest infected cells with accessible provirus, irrespective of reservoir distribution, are poised for reactivation during ART treatment. Together, our findings reveal the extensive inter- and intrapersonal cellular heterogeneity of the HIV reservoir, and establish an initial multiomic atlas to develop targeted reservoir elimination strategies.

## Introduction

The establishment and persistence of the HIV reservoir remains one of the greatest barriers preventing a functional cure in people living with HIV (PLWH). This reservoir, primarily composed of infected long-lived memory CD4+ T cells (Chun et al., 1995, 1997; Finzi et al., 1997), resides in blood and tissue compartments throughout the body and almost invariably reactivates upon antiretroviral therapy (ART) interruption (Deleage et al., 2016; Estes et al., 2017; Pantaleo et al., 1993; Wong et al., 1997). Reservoir persistence results from the stable integration of viral DNA into the genome of a host cell, enabling infection for the lifetime of the cell and its progeny. Substantial strides have been made towards understanding the virological aspects of the HIV reservoir, including intactness, reactivation potential, and integration site landscapes (De Scheerder et al., 2019; Ho et al., 2013). However, beyond population level characterization and quantification, the cellular identity of the HIV reservoir *in vivo* remains an enigma. Defining a surface marker and or epigenetic signature of the cellular HIV reservoir for targeting is of central importance to the HIV cure agenda.

The major impediment towards our understanding of the cellular reservoir is the challenge of identifying infected cells. On average during ART, fewer than 1/1000 CD4+ T cells are infected with HIV in the blood (Eriksson et al., 2013), the majority of which have little to no viral RNA production (Einkauf et al., 2022; McManus et al., 2019). To overcome this, the field to date has used *in vitro* models (Bradley et al., 2018; Golumbeanu et al., 2018), exogenous stimulation strategies (Cohn et al., 2018; Liu et al., 2020), and marker-targeted analysis (many studies including (Fromentin et al., 2016; Neidleman et al., 2020; Pardons et al., 2019)) to identify surface protein and/or transcriptomic properties of the HIV reservoir. These studies have collectively made inroads into the identity of infected cells but have been inherently limited by the experimental systems employed. These limitations, including 1) the heterogeneity of infected CD4+ T cells within bulk populations, 2) incomplete representation of the reservoir by sorting for viral RNA+ and/or protein-expressing cells, and 3) activation-induced malleability of both the cell surface proteome and underlying transcriptome, have obfuscated the precise characterization of infected cells and highlight the need to develop novel strategies to characterize the cellular HIV reservoir at steady state and ultimately following cure-directed strategies.

Here, we used the presence of integrated provirus as a molecular tag to directly identify individual infected cells using single-cell Assay for Transposase Accessible Chromatin (scATACseq) (Buenrostro et al., 2013) and employing a human-viral alignment strategy first developed in a chimeric antigen receptor (CAR) T-cell model (Wang et al., 2020). Further, scATACseq provides unbiased, single-cell epigenetic profiles and can therefore assign cellular identity to cells with and without detectable provirus. An additional goal of reservoir research is to identify cell surface proteins that identify or substantially enrich for infected cells. We therefore combined scATACseq with cell surface protein identity using the recently described ATAC with Select Antigen Profiling by sequencing (ASAPseq) (Mimitou et al., 2021; Swanson et al., 2021). We employed ASAPseq to study both *in vitro* and direct *ex vivo* HIV infection, including the most clinically relevant setting with peripheral blood memory CD4+ T cells from ART-treated PLWH. Our findings provide the first direct and unbiased identification of experimentally and *in vivo* infected memory CD4+ T cells at single-cell resolution with cellular identity. With our multidimensional dataset, we highlight the heterogeneity of HIV+ cells and the variation of surface and epigenetic markers between and across infection contexts. Together, our extensive characterization of HIV infection using ASAPseq results in a multimodal single-cell dataset of cell surface protein composition, epigenetic landscape, and potential regulatory mechanisms to accelerate our understanding of the HIV reservoir for targeted therapy.

## Results

### ASAPseq identification of HIV-infected cells *in vitro*

To determine the potential for ASAPseq to detect HIV-infected cells, we first tested an *in vitro* primary CD4+ T-cell infection model (**Table 1**). CD4+ T cells from an HIV-uninfected individual were activated with anti-CD3, anti CD28/CD49d, and IL-2 for two days, infected with HIV-1 (molecular clone; SUMA), and rested for four subsequent days. We next performed ASAPseq, obtaining 7095 cells that passed quality checks for both ATAC and the antibody-derived tag (ADT) components (**Supplemental Table 1** and **Supplemental Figure 1A**). Following alignment of sequencing data to both human and viral genomes, 1323 cells (18.6%) contained an integrated HIV genome in open chromatin (HIV+, **Figure 1A-B)**. As expected, a smaller proportion of the *in vitro* infected cells expressed p24 protein by flow cytometry (5.1%, **Supplemental Figure 1B**), consistent with known per-cell discrepancies between HIV integration and p24 production (Shan et al., 2017). To confirm the specificity of provirus detection, we aligned an unrelated ASAPseq dataset of uninfected peripheral blood mononuclear cells (Mimitou et al., 2021) using our chimeric human-virus genomes (human (hg38) + SUMA HIV and hg38 + HXB2 HIV). No cells contained reads aligning to either HIV genome, demonstrating the high specificity for ASAPseq-based detection of HIV+ cells (**Supplemental Figure 1C**). Within the HIV+ CD4+ T cells, proviral reads spanned the viral genome but were most prevalent in the long-terminal repeats (LTR) as well as *gag* and *env* genes (**Supplemental Figure 1D**). Fewer reads spanned the *pol* gene consistent with previous findings (Jefferys et al., 2021). Twenty-eight percent of HIV fragments were in LTR regions, 0.01% spanned both LTR and internal (i.e. not LTR), and the remaining fragments (71%) were internal.

**Figure 1:**
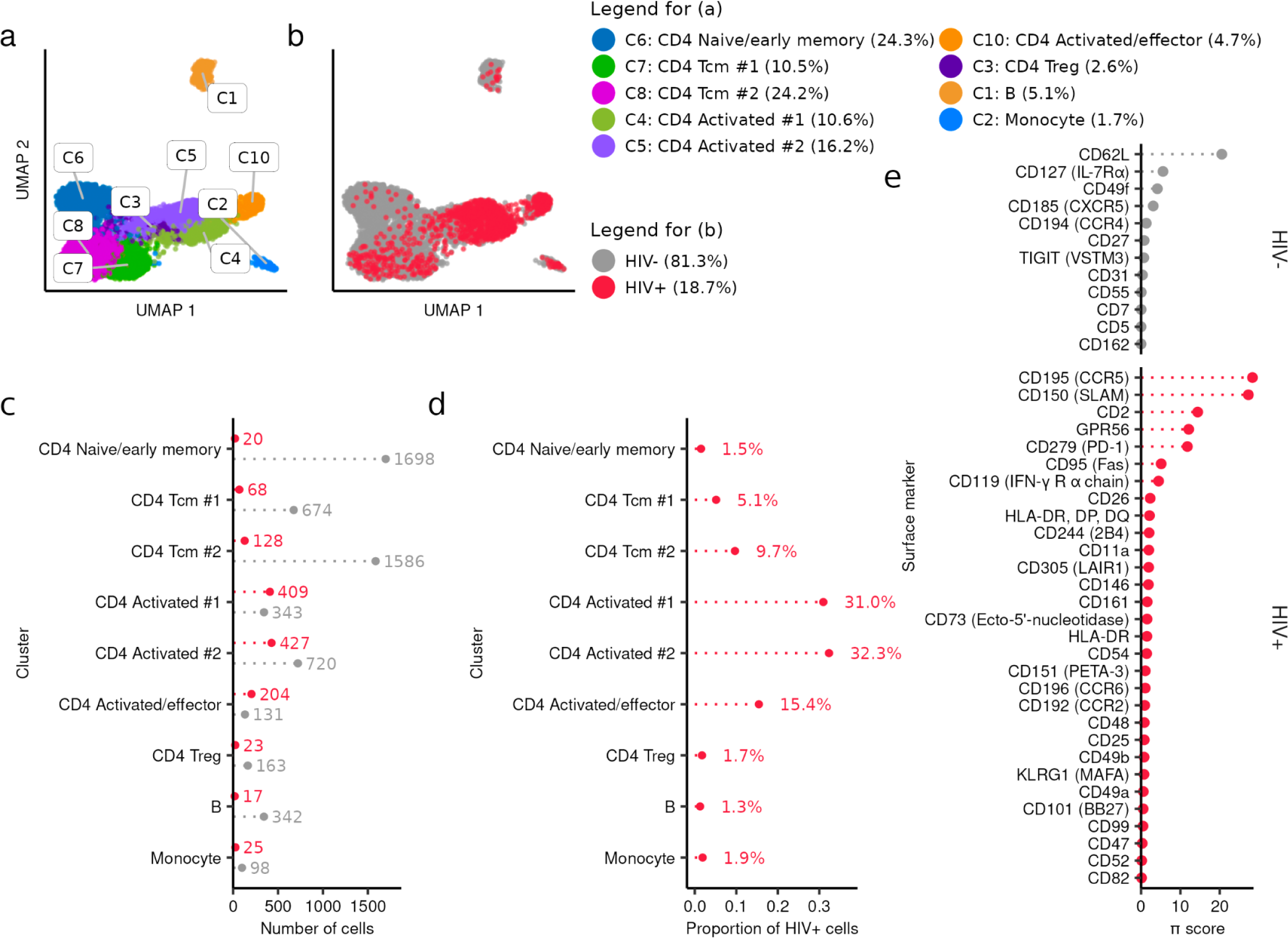
ASAPseq identification of HIV-infected cells in vitro. (A) UMAP representation of ATAC component colored by manually annotated cell phenotypes. (B) UMAP representation of ATAC component colored by detection of HIV reads. (C) Absolute count of cell numbers based on annotated clusters. Red color denotes HIV+ cells while gray color denotes HIV- cells. (D) Percentage of HIV+ cells found in each annotated cluster. (E) Surface enrichment profile in activated HIV+ cells (red) versus activated HIV- cells (gray). Activated cells are defined as the combination of CD4 Activated/effector, CD4 Activated #1, and CD4 Activated #2 clusters. π-score is defined as (-log10(FDR) * log2(FC)).

**Table 1:** Donor information

| Stage | Donor | Sex | Age (y) | Tissues profiled** | Plasma viral load (log copies /ml) |
| --- | --- | --- | --- | --- | --- |
| *in vitro* infection | ND492 | M | 37 | PB | - |
| ART-treated infection | A01 | M | * | PB | <1.7 |
|  | A08 | M | * | PB | <1.7 |
|  | A09 | M | * | PB | <1.7 |
|  | B45 | M | 43 | PB | <1.7 |

\* Age was not reported at the participant level for the original study. However, the age range was reported as between 34 to 52 years.
\*\* PB = peripheral blood

We next examined the phenotypic identity of HIV+ versus HIV- cells. We clustered the cells using the epigenetic data component and annotated based on imputed gene scores from chromatin accessibility and the ADT component (**Figure 1A**, **Supplemental Figures 1E, Supplemental Figures 2, 3,** and **Supplemental Table 2**). Given that PBMCs were enriched for CD4+ T cells prior to infection, most clusters consisted of CD4+ T cells with some contaminating B and antigen presenting (APC) cells. Clusters were separated by activated versus naive/early memory phenotypes, with the greatest number of HIV+ cells (78.6% of all HIV+ cells, **Figures 1B-D**) found within activated and/or effector memory CD4+ T cell clusters. We also observed some HIV+ cells in other CD4+ T-cell phenotypes (18.3%) and a small proportion in monocytes (1.8%). These findings reflect the known phenomenon of preferential *in vitro* HIV infection of activated CD4+ T cells and demonstrate that this feature can be resolved on a single-cell level by ASAPseq.

**Figure 2:**
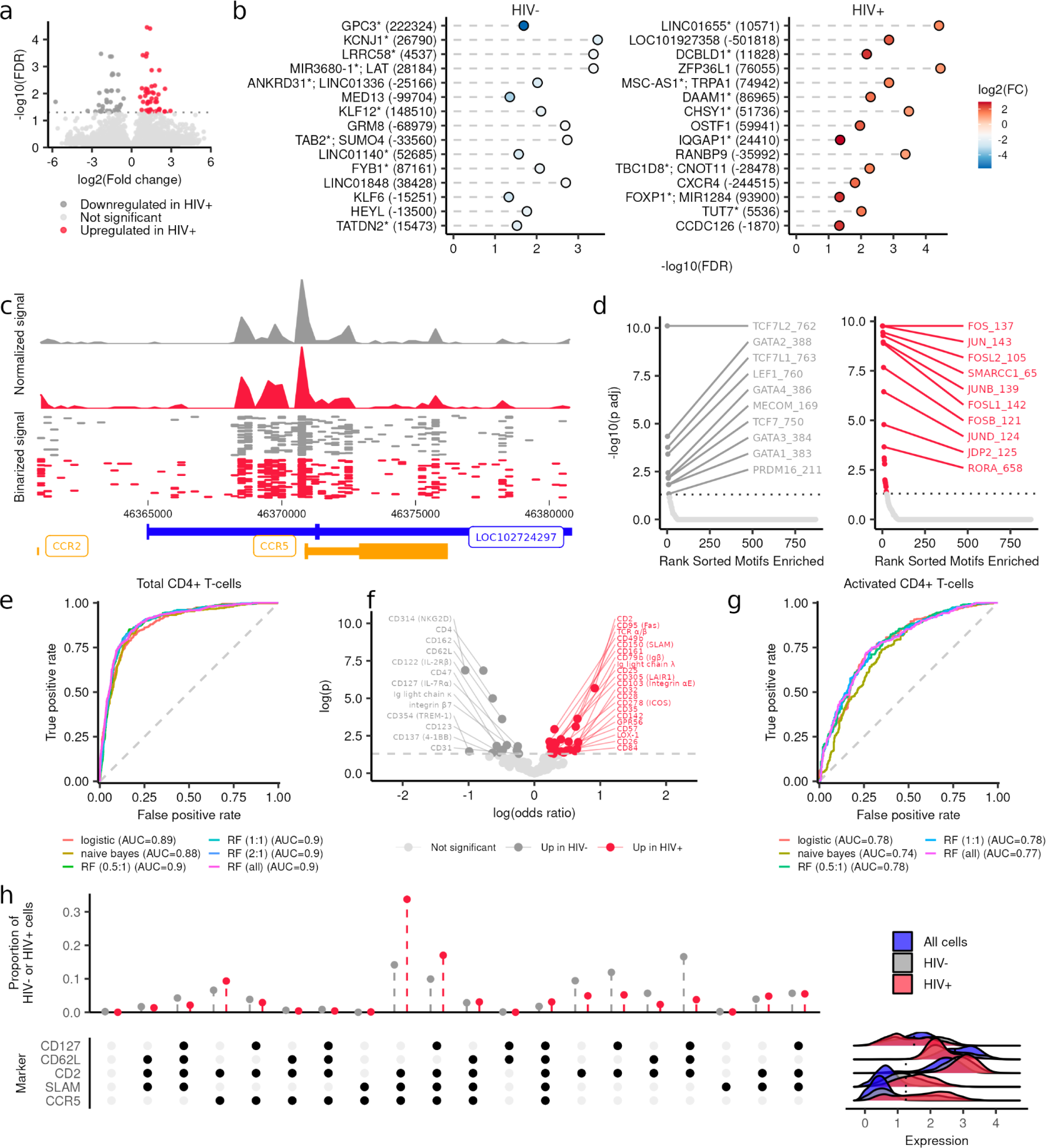
Differential chromatin accessibility in HIV-infected cells and surfacer marker based supervised machine learning in vitro. (A) Volcano plot showing differentially accessible peaks between activated HIV+ versus activated HIV- cells. (B) Top 15 significant peaks in activated HIV- versus activated HIV+ cells. Peaks are ranked by π-score (-log10(FDR) * log2(FC)). Y-axis labels denote nearest TSS and/or if the peak is in a gene (marked by asterisk *) and numbers in parentheses indicate distance to the nearest TSS. Negative numbers indicate that the nearest TSS is upstream of the peak. (C) (top) Aggregated and normalized ATAC signal between activated HIV- (gray) and activated HIV+ cells (red) at the *CCR5* locus. (middle) Binarized ATAC signal at a single-cell resolution (showing random 750 cells for each group). (bottom) Gene map for the genomic region shown: chr3:46360853-46380854 (centered on *CCR5*). Orange genes are in positive orientation while blue genes are in negative orientation. (D) Significant motif enrichments found in the differential peaks (ordered by significance) are shown in activated HIV- cells or activated HIV+ cells. (E) Area under the receiver operating curve (ROC) (AUC) plots for multiple supervised machine learning models for all CD4+ T cells. RF = random forest. Ratios for different RF models indicate the number of HIV- cells used for each HIV+ cell (i.e. 5:1 ratio meant that the HIV- cells in the training dataset were randomly downsampled to get only 5 times the number of HIV+ cells in the training dataset). (F) Significant coefficients for the logistic regression model shown in (E). Positive odds ratio (red) indicates a marker has more weight for HIV+ cells while negative odds ratio (gray) indicates a marker has more weight for HIV- cells. (G) AUC plots for multiple supervised machine learning models for activated CD4+ T cells. (H) Proportion of HIV- (gray) and HIV+ (red) activated CD4+ T cells with different surface marker combinations. In the bottom dot plot, a black dot indicates that the combination is positive for that marker, which was determined by the thresholds indicated with the dotted line on the ridge plots. These ridge plots display the expression distribution of activated HIV+ CD4+ T cells (red), activated HIV- CD4+ T cells (gray), and total cells in the *in vitro* dataset (blue) to help with gating.

**Figure 3:**
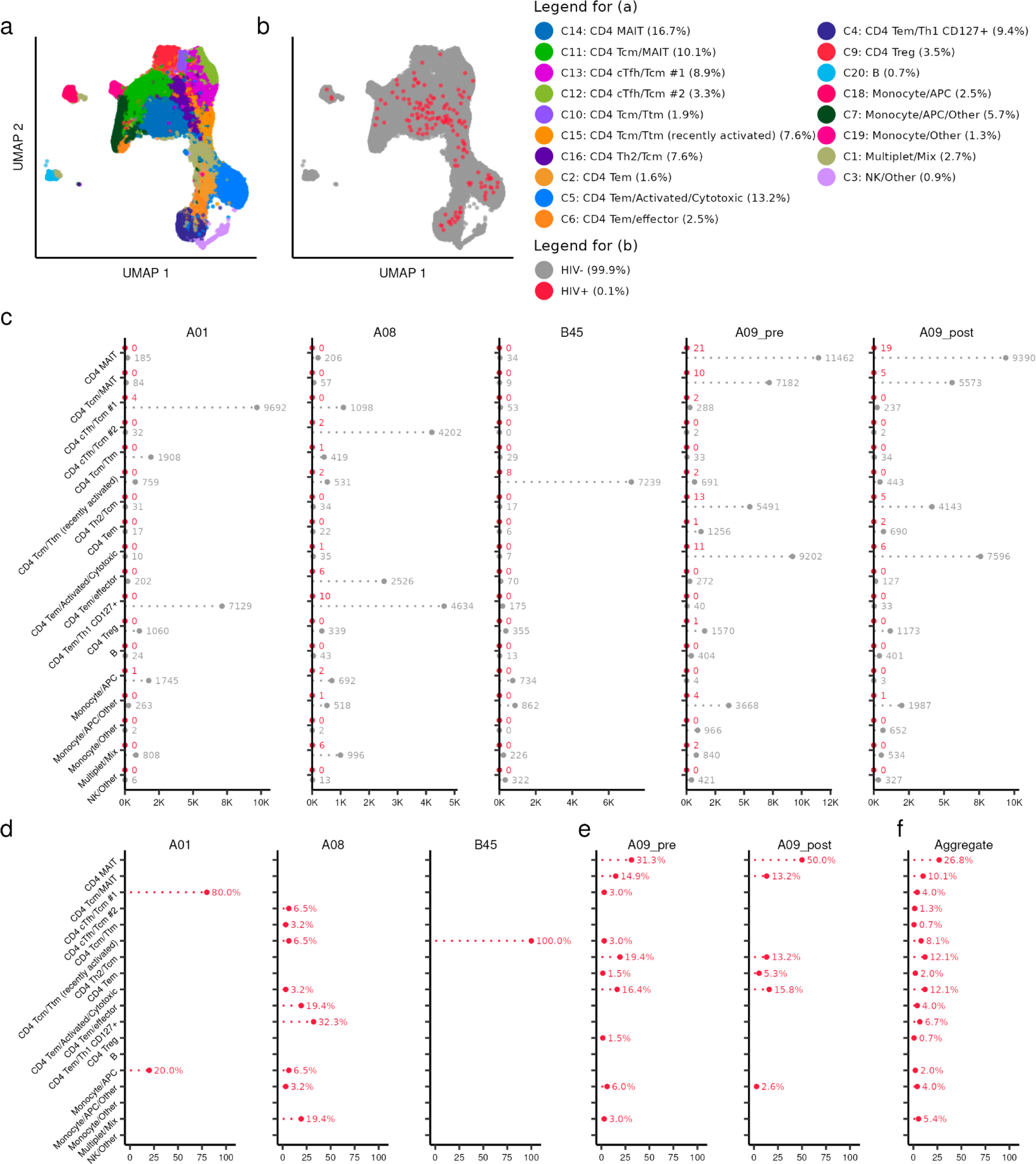
ASAPseg identification of heterogeneous **H**IV-infected cells from peripheral blood of ART suppressed PLWH. (A) UMAP representation of ATAC component colored by manually annotated cell phenotypes. (B) UMAP representation of ATAC component colored by detection of **H**IV reads. (C) Absolute count of cell numbers based on annotated clusters. Red color denotes **H**IV+ cells while gray color denotes **H**IV- cells. (D) Percentage of **H**IV+ cells found in each annotated cluster separated by donors A01, A08, and 845. Y-axis represents percent of the donor’s **H**IV+ cells found in each annotated cluster. **(E)** Percentage of HIV+ cells found before and after treatment interruption in donor A09. **(F)** Percentage of HIV+ cells found in each annotated cluster, aggregated across all ART-treated donors.

In a comparison between all HIV+ and uninfected T cells, we observed significantly higher expression of many surface markers including those associated with T-cell activation on HIV+ cells (**Supplemental Figure 4A** and **Supplemental Table 3**). The top 5 upregulated markers by π-score (Xiao et al., 2014) on HIV+ cells included CCR5, SLAM, PD-1, CD49a, and CD161 while the top 5 upregulated markers on HIV- cells were CD31, CD62L, CD55, CD27, and CD7 (**Supplemental Figure 4A**). The significant upregulation of CCR5 in HIV+ cells is consistent with the usage of CCR5 as the co-receptor for HIV-1 infection (Alkhatib et al., 1996). The other differentially expressed surface markers in HIV+ cells reflect the known preferential infection of activated CD4+ T cells (Zack et al., 1990; Zhang et al., 1999). We therefore asked whether any surface protein distinguished between infected and uninfected cells in the activated clusters (**Figure 1E**, **Supplemental Figure 5**, and **Supplemental Table 3**). In activated HIV+ cells, we found that CCR5, SLAM, CD2, GPR56, PD-1 were the top upregulated markers, while CD62L, CD127, CD49f, CXCR5, and CCR4 were increased in activated HIV- CD4+ cells. This finding suggests that even within the activated/effector cells, the HIV+ cells have higher expression of CCR5 and other activation markers. In the early differentiated HIV+ T cells, we found several significantly increased expression of ICOS, CXCR3, CD25, CD11a, and CD49b while the early differentiated HIV- cells were enriched in CD62L, CD55, CD127, CD162, and CD45RA (**Supplemental Figure 4B** and **Supplemental Table 3**). These surface antigen marker findings highlight the heterogeneity of infected cells even in the activated cell population.

### Differential chromatin accessibility in HIV-infected cells and surfacer marker based supervised machine learning *in vitro*

We next examined whether there was any differential genomic accessibility between activated HIV+ and activated HIV- cells. We observed heightened accessibility in several genomic regions in activated HIV+ cells (**Figures 2A-B** and **Supplemental Table 4**). Several nearest genes to these peaks have been previously implicated for HIV infection and replication. We also observed a greater accessibility at a peak ∼1kb upstream of the *CCR5* transcription start site (**Figure 2C**), thus showing a potential concordance between CCR5 surface expression and gene accessibility. From these differentially accessible genomic regions, we assessed transcription factor motif enrichment. In the activated HIV- cells (**Figure 2D**), the top enriched motifs include several members of the TCF family including TCF7 (TCF-1) and LEF1, both regulators of CD4+ T-cell development and differentiation (reviewed in (Ma et al., 2012)). In the activated HIV+ cells (**Figure 2D**), the top motifs included proliferation and activation motifs in the AP-1 related family such as the JUN and FOS families (reviewed in (Foletta et al., 1998)). These findings, in combination with the surface marker profiles, are concordant with the known immunobiology of HIV-1 infection, thus showing that ASAPseq is a viable technique for ascertaining phenotypes of HIV+ cells.

Finally, as an initial step towards selectively recognizing and targeting HIV+ cells, we employed several supervised machine learning methods to determine whether any cell surface proteins could, in combination, predict infected cells. We assessed logistic regression, naive Bayes, and random forest methods using a 70/30 (training/test) split of our dataset. Using all CD4+ T cells, we found similar results between the models with area under the receiver operating curve (ROC) (AUC) values of 0.89, 0.88, and 0.9 respectively (**Figure 2E**) suggesting a reasonable classification of HIV+ cells *in vitro* with this current ADT panel. In the logistic regression, markers with the greatest odds ratio and significance included CD2, CD49b, CD95, SLAM, and TCR α/β for classifying HIV+ cells while CD4, NKG2D, CD62L, and CD162 were the top markers for classifying HIV- cells (**Figure 2F**). For the random forest model, the top markers by importance (measured as mean decrease in accuracy when permuted) were CCR5, SLAM, CD49b, CD2, and CD95. When applied to only activated CD4+ T cells, the random forest model and logistic regression both performed the best with an AUC of 0.78, compared to 0.74 with naive Bayes. Compared to the total CD4+ T-cell models, there were higher false positive rates suggesting that additional markers tailored to activated T cells might benefit classification efforts (**Figure 2G**). The activated CD4+ T-cell random forest was driven primarily by CCR5, SLAM, CD69, CD2, and GPR56 in decreasing order of importance. Overall, these supervised machine learning models are in concordance with our differential expression testing and highlight the prominent roles of CCR5, SLAM, and CD2 as markers of HIV-1 *in vitro* infection (**Figure 2H**).

### ASAPseq identification of heterogeneous HIV-infected cells from peripheral blood of ART suppressed PLWH

We next applied ASAPseq to blood CD4+ T cells in the setting of ART to assess the reservoir in the most accessible compartment and in a clinically relevant context. To define the characteristics of the HIV CD4+ T-cell reservoir in the blood of ART-treated PLWH (ART-PLWH), we purified memory CD4+ T cells from the PBMC of 4 ART-PLWH and performed ASAPseq. Given the rarity of infected cells during ART treatment (Chun et al., 1997) and the relative paucity of infected cells in peripheral blood compared to tissues, we substantially increased the number of memory CD4+ T cells assessed compared to our in vitro infection studies. Altogether, the combined ART-treated dataset contained 127761 cells of which 149 (0.12%) cells were HIV+ (**Supplementary Figure 6A** and **Table 2**). For all donors, we aligned the ATAC reads to autologous HIV sequences (near full length or HIV *env*) and HXB2 to increase the odds of detecting infected cells due to the prevalence of proviral mutations in long-term PLWH (Ho et al., 2013). With the autologous alignments, most aligned regions were primarily detected in the LTR and/or *nef* region (**Supplementary Figure 7**). Cells from donor A09 during both timepoints also had internal fragments that mapped to *gag*, *pol*, and *env* genes (**Supplementary Figure 7**).

**Table 2:** Cell counts stratified by individual

| Individual | Total cells | HIV+ cells<br>(% of total cells) | Total HIV DNA copies per<br>million CD4+ T cells<br>(percentage)<br>(from Salantes et al., 2018) |
| --- | --- | --- | --- |
| A01 | 23962 | 5 (0.02%) | 293.8 (0.029%) |
| A08 | 16398 | 31 (0.19%) | 1564.5 (0.16%) |
| A09 | 77242 | 105 (0.14%) | 1297.3 (0.12% pre-ATI)<br>1221.8 (0.12% post-ATI) |
| B45 | 10159 | 8 (0.08%) | - |

After clustering based on the ATAC component and annotating with both ADT and ATAC components (**Supplementary Figure 6B**, **Supplementary Figures 8-9**, and **Supplemental Table 6**), we identified multiple memory CD4+ T cell subsets with only small numbers of contaminating cells (**Figures 3A-B**). The T cell clusters separated based on epigenetic profiles associated with T-cell differentiation states. For example, central memory T cells (Tcm) had distinct epigenetic and surface antigen profiles compared to those seen in effector/effector memory (Tem) cells (**Supplementary Figures 8-9**). Across the ART-PWLH, we observed no consistent cell type that was predominantly infected. Some donors (A01 and B45) showed more dominant populations of HIV+ cells within a single cell type while other donors (A08 and A09) had more detections across different cell types (**Figure 3C**). In sharp contrast with the heterogeneity across donors, we noticed a remarkable maintenance of cell types amongst the HIV+ cells between the pre- and post-ATI timepoints from donor A09 (**Figure 3D**), indicating relative compositional stability of the cellular phenotypes of the HIV reservoir in this donor after ATI. After aggregating all HIV+ cells across donors, we detected HIV+ cells within multiple subsets including CD4+ Tcm, Th2, circulating Tfh (cTfh), mucosal associated invariant T (MAIT) cells and Tem/effector cells (**Figures 3B,E**). We also detected a small proportion of HIV+ cells within the contaminating APC clusters, but it is unclear whether these were in actual APCs, CD4+ T cells included in doublets with APCs, or miscalled as APCs.

### High degree of heterogeneity in HIV-infected cells from peripheral blood of ART suppressed PLWH

We next assessed whether differential surface markers were present between the HIV+ CD4+ T cells versus the HIV- CD4+ T cells. Using the statistical test employed by Seurat and DESeq2, we found significant enrichment of CD99 and CD2 in the HIV+ cells, markers associated with T cell adhesion and co-stimulation, respectively. We found enrichment of several markers in the HIV- T cells including CD200, CD74, CD41, and NKG2D (**Figure 4A** and **Supplemental Table 6**). With another statistical test (Wilcoxon), we found enrichment of different markers including activation markers such as SLAM, CCR2, CD49d, CD26, HLA-DR, and PD-1 that were more highly expressed on HIV+ CD4+ T cells (**Figure 4B**, **Supplemental Figure 10**, and **Supplemental Table 6**). Similar to the results from the DESeq2 method, we found that CD200 and CD74 were also more expressed on HIV- CD4+ T cells. It is important to note that the enrichment of some of these markers (from either test) can vary across donors as seen with PD-1 primarily being driven by individual A08 (**Supplemental Figure 10**).

**Figure 4:**
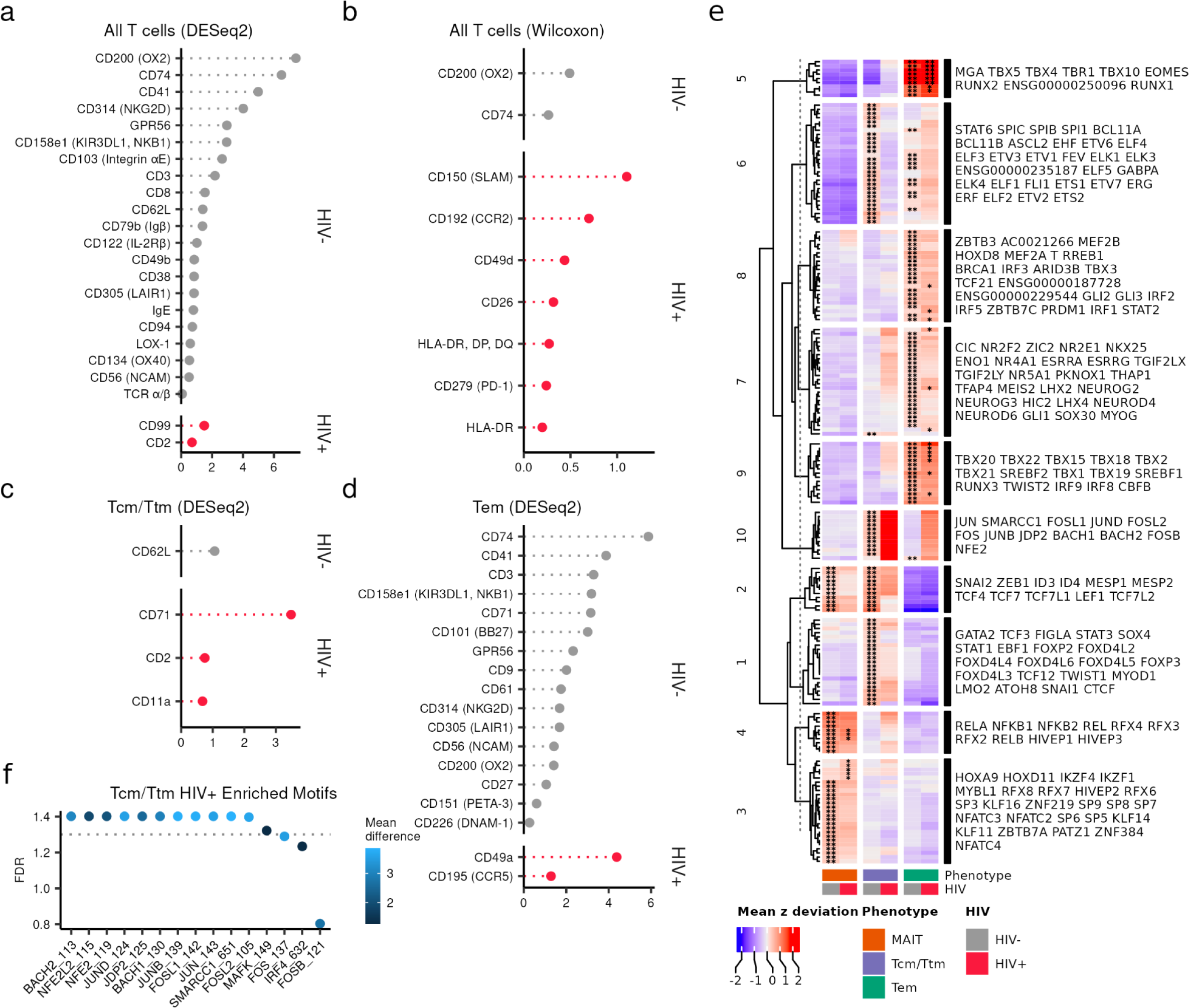
High degree of heterogeneity in HIV-infected cells from peripheral blood of ART suppressed PLWH. (A) Surface enrichment profile in all HIV+ CD4+ T cells (red) versus HIV- CD4+ T cells (gray). Test performed using the DESeq2 pseudobulk method in Seurat. (B) Surface enrichment profile of the same cells as in (A) but using the Wilcoxon statistical test in Seurat. (C) Surface enrichment profile of Tcm/Ttm HIV+ (red) versus Tcm/Ttm HIV- cells (gray) using the DESeq2 method. Tcm/Ttm cells are defined by the combination of all clusters containing the terms “Tcm” or “Ttm” (but not MAIT) in Figure 3A. (D) Surface enrichment profile of Tem HIV+ (red) versus Tem HIV- (gray) cells using the DESeq2 method. Tem cells are defined by the combination of all clusters containing the term “Tem” in Figure 3A. X-axis units for (A-D) are in π-score (see Figure 1E legend). Heatmap of mean chromVAR deviations (z-score) of transcription factor motifs from CISBP database (displayed by row) that define each aggregate group (MAIT, Tcm/Ttm, and Tem) separated by HIV- versus HIV+ (displayed by column). Motifs were selected by FDR < 0.05 and a mean difference > 0.5, indicating a significant accessibility of regions containing a given transcription factor motif for cells in the cluster (indicated by asterisks) as compared to all other cells (getMarkerFeatures function in ArchR). Each cell is colored by the mean deviation (z-score) where values > 0 indicate positive enrichment of a motif. Motifs were clustered using k-means clustering and the motifs are labeled to the right of the heatmap in order from top to bottom for each cluster. Asterisks indicate that the motif was significantly enriched in the specific group (column). * < 0.05; ** < 0.01 (F) chromVAR deviations were compared between HIV+ Tcm/Ttm cells and HIV- Tcm/Ttm cells and the top 15 most significant motifs are shown. The dotted line indicates a -log10 transformed FDR value of 0.05. Color indicates the mean difference in chromVAR deviations (value > 0 indicates enrichment in HIV+ cells).

To reduce the variance caused by differential expression of markers based on memory and functional differentiation (i.e. central/transitional memory, effector memory, and MAIT), we conducted differential expression testing between HIV+ and HIV- cells in the following aggregated CD4+ T cell groups: Tcm/Ttm cells (**Figure 4C**), Tem cells (**Figure 4D**), and MAIT cells. Other cell types were excluded for this analysis due to low HIV+ cell counts. In the Tcm/Ttm cells, we found enrichment of CD71, CD2, and CD11a on HIV+ cells while there was an enrichment of CD62L on the HIV- cells indicating that the Tcm/Ttm cells were contributing to the CD2 from the bulk CD4+ T cell analyses (**Figure 4C**). In the Tem cells, the integrin CD49a and receptor CCR5 were significantly enriched on HIV+ cells, while multiple markers including CD74, CD41, and CD3 were enriched on HIV- Tem cells (**Figure 4D**). No significant markers were differentially expressed for either HIV+ or HIV- MAIT cells (data not shown). These results highlight the surface marker and phenotypic heterogeneity present in specific memory phenotypes as well as the inability of any marker amongst the 154 antibodies tested to uniquely identify HIV+ cells.

To address this heterogeneity in a more unbiased way, we next assessed if there were unique epigenetic signatures between the HIV+ and HIV- CD4+ T cells. We used chromVAR, specifically designed for differential motif analysis from sparse scATACseq data (Schep et al., 2017), on all HIV+ versus HIV- cells grouped by the same three aggregate memory/functional phenotypes (Tcm/Ttm, Tem, and MAIT) in the surface marker analyses. We identified several highly enriched motifs in the different groups over all other cells that are not in the analyzed group (collectively referred to as background cells). Motifs were clustered into different modules via hierarchical clustering (**Figure 4E**). In support of our manual annotation, Tcm/Ttm cells showed greater enrichment of Tcf7 and Lef1 motifs (module 1) compared to Tem cells, while the Tem cells showed greater enrichment of Tbx21 (T-bet), Eomes, and interferon related motifs (modules 5 and 9). MAIT cells, which can be heterogeneous in terms of different memory phenotypes, showed enrichment in motifs from the nuclear factor-kappaB (NF-κB) transcription factor family including NFKB1, REL, RELA, and RELB (modules 3 and 4).

While HIV+ and HIV- cells shared similar motifs within the aggregate clusters, we observed patterns of greater enrichment of certain motifs in the HIV+ cells. Module 10 - consisting of BACH2 and AP-1 related family transcription factors such as Jun and Fos - was highly enriched for HIV+ Tcm/Ttm and Tem cells, but was not significant when compared to the background of all other cells. When comparing Tcm/Ttm HIV+ versus Tcm/Ttm HIV- cells, we found significant enrichment of motifs associated with module 10 (**Figure 4F**). HIV+ MAIT cells curiously did not show an enrichment in this module but did show significant enrichment in HOXA9, HOXD11, IKZF4, IZKF1, and MYBL1 motifs over background cells (**Figure 4E**). Altogether, these motif signatures across our dataset indicate that Tcm/Ttm and Tem HIV+ cells share an epigenetic signature consistent with a heightened immune activated state.

## Discussion

The targeted elimination of the HIV reservoir requires a deep understanding of both the virologic and cellular characteristics. Prior work has yielded extensive knowledge of viral diversity, integration, clonal expansion, intactness, and latency dynamics of the viral reservoir, allowing a deeper understanding of the target for clearance. However, precise definition of cellular reservoir has proven more difficult due to the rarity of infected cells and the challenge of identifying infected cells while preserving their native state. To address this, we applied a novel single-cell genomic and bioinformatic strategy using ASAPseq to identify infected cells by the presence of integrated proviral DNA in the multiomic context of single cell epigenetic and surface antigen profiling. Using this strategy, we found that cells with proviral DNA can indeed be directly detected and individually characterized directly from the peripheral blood of PLWH on ART. We anticipate that this strategy will have broad application for in depth analysis of the HIV reservoir within different cell populations, therapeutic interventions, and HIV cure and eradication studies (Abdel-Mohsen et al., 2020).

Numerous studies have attempted to identify cell surface markers and memory phenotypes that enrich or specifically identify CD4+ T cells harboring integrated proviral DNA. Recent studies have shown that Tem cells contain high levels of transcriptionally active proviruses (Collora et al., 2022; Grau-Expósito et al., 2019). As such, we expected Tem cells to be enriched in our dataset due to the assumed accessible chromatin. However, we find no Tem bias in the cells harboring integrated virus as HIV+ cells in ART-PLWH from our study were distributed amongst a wide spectrum of memory phenotypes. Importantly, this distribution also differed between individuals but not within individual A09 where we observed similar phenotypes of infected cells that were conserved before and after ATI. This indicates that ATI does not drastically shift the reservoir and is observed along with the reported stability of the reservoir with ATI (Salantes et al., 2018). The lack of memory bias in our dataset might be attributed to the differences in when ART was initiated during infection and the duration of ART, which are factors that can impact reservoir dynamics (Chéret et al., 2015; Chun et al., 2007; Massanella et al., 2021). Until now, this degree of heterogeneity across individuals and longitudinal conservation of phenotypes across ATI has not been demonstrated and indicates the necessity of single-cell based approaches.

Our surface antigen profiling assessed many individual markers previously associated with HIV infection, including PD-1 (Perreau et al., 2013), CXCR5 (Perreau et al., 2013), CCR6 (Gosselin et al., 2017), CTLA-4 (McGary et al., 2017), CD69 (Cantero-Pérez et al., 2019), OX40 (Kuo et al., 2018), CD2 (Iglesias-Ussel et al., 2013), Lag-3 (Pardons et al., 2019), TIGIT (Fromentin et al., 2016; Pardons et al., 2019), CD20 (Serra-Peinado et al., 2019), and CD161 (Li et al., 2019), but did not include CD30 (Hogan et al., 2018), α4β1 (Pardons et al., 2019), CD32a (Abdel-Mohsen et al., 2018; Descours et al., 2017), or Sialyl-LewisX (Colomb et al., 2020). Importantly, between the different modalities we tested, no single previously identified cell surface marker was solely associated or predictive of the presence of integrated provirus. We do, however, find that specific combinations of markers, both by presence and relative level, may more reliably enrich for HIV-infected cells on *in vitro* infected samples. For example, CCR5 and PD-1 were significantly expressed on HIV+ over HIV- cells for bulk and activated T cells i*n vitro* and Tem cells *ex vivo* from ART-treated peripheral blood, which represent known associations with the HIV reservoir (Fromentin et al., 2016; Perreau et al., 2013). However, CCR5 was not heightened on HIV+ cells in Tcm/Ttm cells from ART-PLWH. We instead observed higher expression of CD71 expression, which correlates with cell cycling (Younes et al., 2018) and is a unique surface marker for ferroptosis (Feng et al., 2020). This could suggest that these infected cells are dividing frequently and are potentially predisposed to cell death. We also observed an enrichment of CD2 which has been highlighted by prior work in enriching for HIV DNA in ART-PLWH (Iglesias-Ussel et al., 2013). The integrin CD11a was also enriched in Tcm/Ttm and is known to be increased upon HIV-1 infection (Rossen et al., 1989).

Identification of cell surface markers found preferentially on HIV uninfected cells is also useful. We found higher levels of the inhibitory receptor CD200 (OX2) that appeared to be driven by Tem cells in ART-PLWH. These may be indicative of HIV-specific cTfh cells which are known to express CD200 and PD-1 more frequently than non-HIV-specific cTfh. For these HIV-specific cTfh cells, their frequency and expression of inhibitory receptors are not modulated during ART treatment (Niessl et al., 2020). We also curiously noticed an increase in CD74, which is the invariant chain for major histocompatibility complex class II (MHC-II) which also appears to be driven in Tem cells. However, the functional significance of decreased expression of CD74 on HIV+ cells is less clear. Our findings, as a whole, indicate that HIV+ cells are highly heterogeneous at interpersonal and intrapersonal levels, and that each cell subset and compartment needs to be evaluated individually. Each of these individual subsets has its unique biology, function, and differentiation potential that will contribute in currently unknown ways to the establishment and maintenance of the reservoir. Along with the variable impact of latency reversal agents on different memory subsets (Pardons et al., 2019), our results highlight the difficulty in finding a strong candidate for a “shock and kill” therapy as the cells harboring proviruses have different epigenetic phenotypes.

The epigenetic modality afforded by our single-cell multimodal strategy also yielded insights into both known and potential differential regulatory factors that impact HIV infection *in vitro* and *ex vivo*. Infection of *in vitro* activated CD4+ T cells was highly associated with increased accessibility upstream of the *CCR5* transcription start site, which appears to be associated with increased CCR5 protein expression. In this same system, we further identified accessible peaks near genes previously implicated in HIV infection and replication, including *DCBLD1, GGA2*, and *PLEKHA3*. *DCBLD1* enhanced HIV replication during a RNA interference screen (Zhou et al., 2008), while *GGA2* is important during downregulation of CD4 by Nef (Landi et al., 2014). Knockdown of *PLEKHA3* was shown to inhibit HIV-1 replication (Brass et al., 2008)

In the case of ART-PLWH, the Tcm/Ttm/Tem HIV+ cells had a high enrichment (though not significant when compared to background cells) of a motif module that consisted of AP-1 related transcription factor motifs and Bach1/Bach2 motifs. The presence of AP-1 motifs suggests that these cells are poised for or have a history of activation/proliferation which may indicate an increased likelihood for viral RNA transcription and reactivation upon ART interruption or treatment with latency reversal agents. Recent reports have suggested that Tem/effector cells are the prominent memory pool with active transcription at the provirus locus during ART (Collora et al., 2022; Einkauf et al., 2022). Given our proclivity for finding accessible proviruses, our data indicates that the epigenetic environment in these cells may be primed for such activity or are actively transcribing the provirus. Furthermore, many motifs in this module were significant in comparing HIV+ versus HIV- Tcm/Ttm cells - thus indicating that even the Tcm/Ttm HIV+ pool may also contain this poised state. We also curiously noticed high (but not significant) enrichment of NFKB1 motifs in MAIT HIV+ cells and Tcm HIV+ cells but not Tem HIV+ cells. *NFKB1* encodes for NF-kB, which is an important pioneering factor that can lead to the chromatin accessibility and a major transcription factor needed for HIV transcription - thus reinforcing the notion of priming for reactivation. These findings along with the impact of other modules will need to be validated with future studies that can incorporate ASAPseq modalities along with transcriptomic analyses to assess HIV viral RNA production. Taken together, our epigenetic analysis provides a framework and new opportunities for understanding the complex regulation of HIV infection and ongoing processes regulating uncontrolled infection.

One major limitation of ASAPseq to identify infected cells is the requirement for the provirus to be in accessible chromatin. This may bias our surface and epigenetic profiling of virally infected cells towards an activated phenotype or selectively profile only the “active” reservoir (Einkauf et al., 2022). Proviruses become less accessible over time *in vitro* (Lindqvist et al., 2020) and after extended ART (du Chéné et al., 2007) which is a target for the cure field through epigenetic modification with latency reversal agents (Darcis et al., 2015; Grau-Expósito et al., 2019; Leth et al., 2016; Reuse et al., 2009). This limitation is inherent in other reservoir measurement techniques (including QVOA and IPDA) which are restrained by incomplete proviral awakening via cellular stimulation or by primers being incompatible with sequence diversity, respectively (Kinloch et al., 2021). Despite this limitation, the detection of only accessible provirus by ASAPseq still offers unparalleled insight into those infected cells capable of reactivation or transcription at the proviral locus. Another limitation is the difficulty in assessing proviral intactness due to short-read sequencing and our transposase-based methodology. Alignments towards multiple proviral genes in a single cell can be used as a proxy for intactness given that most defective proviruses have large deletions in treated PLWH (Ho et al., 2013). In the *in vitro* infected cells, we found broad coverage of reads to the HIV genome and a large detection of infected cells, including those not actively expressing p24 protein. The *ex vivo* infected cells predictably yielded much sparser proviral sequence coverage. We also observed a dropoff in sequence recovery from *pol* regions, possibly explained by nucleosomal occupancy (Rafati et al., 2011) or more persistent transcription factor binding (Van Lint et al., 1994). The inclusion of scRNAseq as a third modality (Mimitou et al., 2021; Swanson et al., 2021) can assist in addressing intactness but is also limited towards accessible proviruses. Our focus on HIV DNA+ cells, regardless of intactness or transcription, is still relevant given that these cells survived clearance by the immune system and/or the short half-life associated with HIV infection kinetics (Perelson et al., 1996). Cells with defective proviruses from treated PWLH are still relevant as viral proteins can still be produced (Duette et al., 2022; Imamichi et al., 2020), which may interface with the immune system.

Overall, we have highlighted the complex heterogeneity of the HIV reservoir by using a novel unbiased genomic strategy to identify HIV infected cells at single-cell resolution with simultaneous surface antigen and epigenetic data. With this unparalleled resolution via ASAPseq compared to currently published strategies for studying the HIV reservoir, we uncovered both known and novel surface markers of HIV infection as well as accessible transcription factor motifs that may regulate or be indicative of HIV infection. Our study paves the way for future unmanipulated studies of the HIV reservoir. Furthermore, this approach can also be applied for lentiviral-based immunotherapy and other latent infections that involve nuclear, viral DNA. Our strategy and data are the start of a multiomic atlas of HIV infected cells with phenotypic and epigenetic characteristics - thus contributing towards the united goal of identifying the HIV reservoir for a targeted, functional cure of HIV.

## Methods

### Study approval

ART-treated samples, A01, A08, and A09, were provided from the ACTG clinical trial A5340, which was conducted with protocols set forth by the Institutional Review Boards at the University of Pennsylvania and the University of Alabama at Birmingham and was previously published (Bar et al., 2016). All donors for this study provided informed consent in compliance with protocols set forth by the respective institutional review boards. Another ART-treated PLWH, B45, was recruited from the BEAT-HIV program cohort where an apheresis sample was collected under ART.

### Samples

For the *in vitro* infection model, peripheral blood mononuclear cells (PBMCs) were obtained from an HIV-negative donor apheresis from the Human Immunology Core (HIC) at the University of Pennsylvania. PBMC were cryopreserved at the time of receipt. For ART-treated studies, peripheral blood (PB) samples were acquired via apheresis and cryopreserved. Samples A01, A08, and A09 had previously experienced therapy interruption through the ACTG A5340 trial, but were under at least 6 months (for the post-ATI timepoint) of ART suppression at the time of analysis. Peripheral blood mononuclear cells were prepared as previously described (Nguyen et al., 2019).

### In vitro infection model

Uninfected PBMC were thawed and rested overnight in a humidified incubator at 37 °C. CD4+ T cells were negatively enriched (STEMCELL Technologies, #19052), pelleted, and resuspended in complete RPMI media at a concentration of 2e6 cells/ml in a 6-well tissue culture plate. An activation cocktail consisting of anti-CD3 (1 μg/mL; Bio-Rad #MCA463EL), recombinant IL-2 (100 U/mL; Sigma Millipore #11011456001), and anti-CD28 + anti-CD49d (1.5ug/ml; BD #347690) were added to the cells. The cells were activated for 2 days in a humidified incubator at 37 °C. Cells were then pelleted in a 15ml tube and resuspended in 1ml of complete RPMI. HIV-1 virus stock (strain SUMA; provided by the University of Pennsylvania CFAR Virus and Reservoirs Core) was thawed briefly in a 37 °C water bath. 50ng (p24) of viral stock was added to the suspension and mixed by pipetting. Cells were then infected by spinoculation for 45 minutes at 400 x g. Cells were rested in a humidified incubator at 37 °C for 1 hour (cap loosened to promote gas exchange) before washing with complete RPMI and pelletting. Cells were resuspended at a concentration of 2e6 cells/ml and left to rest for 2 days in a 6-well tissue culture plate. Complete RPMI was added to the cells after two days. The cells were collected after an additional two days and pelleted for dead cell removal based on Annexin V using the Dead Cell Removal (Annexin V) Kit (STEMCELL Technologies, #17899).

### Flow cytometric verification of HIV-1 infection

Staining and flow cytometry were based on a previously published protocol (Kuri-Cervantes et al., 2020). Approximately 1.5 million cells from the *in vitro* infection culture were spun down at 400 x g for 5 min and resuspended in 45ul of PBS. Live/dead staining was performed using 5μl of a 1:60 dilution stock of prepared Live/Dead Fixable Aqua Dead Cell Stain (Invitrogen). Cells were stained for 5 minutes in the dark at room temperature. A staining cocktail with FACS buffer and CD8 BV570 (BioLegend #301038, clone RPA-T8) was added for a 10 minute stain in the dark at room temperature. 1ml FACS buffer was added and the cells were spun down at 400 x g for 5 min. Cells were permeabilized with 250μl of BD Cytofix/Cytoperm solution (BD #554714) for 18 minutes in the dark at room temperature.

1ml of BD Perm/Wash Buffer (BD #554714) was added and cells were spun down at 600 x g for 5 min. After supernatant was discarded, cells were resuspended in staining solution containing anti-p24 FITC (Beckman Coulter #6604665; clone KC57) and BD Perm/Wash Buffer for a final staining volume of 50μl. Cells were stained in the dark for 1 hour at room temperature. Cells were washed with 1ml of BD Perm/Wash Buffer and fixed with 350ul of 1% paraformaldehyde. Data were acquired on a BD FACS Symphony A5 cytometer and analyzed using FlowJo (Version 10, BD).

### Memory CD4^+^ T-cell enrichment

Memory CD4+ T cells were enriched by negative selection using the EasySep Human Memory CD4+ T Cell Enrichment Kit (STEMCELL Technologies #19157) following the recommended protocol. After collecting enriched memory CD4+ T cells, the cells were spun down at 400 x g for 5 min and resuspended in 500μl PBS. Cells were counted with trypan blue staining using a Countess II (Invitrogen) prior to beginning the ASAPseq protocol.

### ASAPseq - cell preparation and staining

Buffer and cell preparation were performed as previously published (Mimitou et al., 2021) and as summarized below. 5e5 to 1e6 cells were resuspended in 22.5μl of Staining Buffer and incubated with 2.5μl of TruStain FcX (BioLegend #422302) for 10 minutes on ice. One test of the TotalSeq-A Human Universal Cocktail, V1.0 (BioLegend #399907) was prepared according to the manufacturer’s protocol with ASAPseq Staining Buffer. 25μl of the antibody cocktail was added to the cells for 30 minutes on ice. Cells were washed with 1ml Staining Buffer and pelleted (all spins at 400g for 5 minutes with centrifuge set at 10C) for 3 total washes. Cells were then resuspended in 450μl of PBS. 30ul of 16% paraformaldehyde was added to fix the cells for 10 minutes at RT with occasional swirling and inversion. The fixation reaction was quenched with 25.26μl of 2.5M glycine solution. Cells were then washed and pelleted with 1ml of ice-cold PBS for a total of two washes. Fixed cells were lysed using 100μl of OMNI lysis buffer for 3 minutes on ice. Cells were then washed and pelleted with 1ml Wash Buffer (spin at 500g for 5 min with centrifuge set at 10C). After supernatant was removed, cells were resuspended in at least 150μl (depending on original cell input) of 1X Nuclei Buffer (10X Genomics) and strained to remove aggregates using a 40μm FlowMi strainer (Sigma #BAH136800040-50EA). Cells were counted using trypan blue staining to verify successful permeabilization (>95% trypan blue positive staining). Cells were diluted as needed according to the scATACseq Chip H protocol (10X Genomics).

### ASAPseq - library preparation

Single-cell droplets were generated using the Chromium platform and a scATACseq Chip H kit (10X Genomics). Modifications mentioned in the original ASAPseq protocol were performed to allow for capture of antibody-derived tags (ADT). Library preparation for the ATAC libraries were performed as recommended by 10X Genomics. ADT libraries were prepared as recommended by the ASAPseq protocol with 3μl of the silane bead elution used as additional input to the ADT indexing PCR.

### ASAPseq - library sequencing

Sequencing runs were performed on NextSeq 550 or NovaSeq 6000 platforms (Illumina) with a target of at least 10,000 reads/cell for ADT libraries and 25,000 reads/cell for ATAC libraries.

### Initial ATAC data processing

Libraries sequenced via the NovaSeq were filtered for potential index hopping by using index-hopping-filter (10X Genomics; v1.1) in ATAC mode. Filtered reads were then processed the same as other libraries sequenced with the NextSeq. A chimeric genome was built using cellranger-atac mkref (10X Genomics; v2.0.0) with the default hg38 genome (10X Genomics; refdata-cellranger-arc-GRCh38-2020-A-2.0.0) along with either SUMA_TF1 (for *in vitro*) or autologous sequences + HXB2 (for ART) (**Supplemental File 1** and **Supplemental Table 7**). The motifs.pfm file from the default hg38 genome reference was copied over to the new chimeric genome to assist with quality control during downstream processing by cellranger. The reads were aligned and counted to their respective chimeric genome using cellranger-atac count. Raw fastq files and processed cellranger-atac files are deposited in the NCBI Gene Expression Omnibus (GEO) under accession number GSE199727. After alignment, barcodes pertaining to multiplets were selected using AMULET (Thibodeau et al., 2021) for downstream filtering. Fragment files from cellranger-atac were loaded into ArchR (v.1.0.2) (Granja et al., 2021) for downstream analysis. Arrow files were created for each sample. Cells were filtered for TSS Enrichment ≧ 8. Barcodes that were selected by AMULET as being multiplets were then filtered out of the Arrow file. Samples, by level, continued in the ArchR pipeline. Briefly, iterative latent semantic indexing, Harmony (for batch effect correction as needed), cluster generation (via Seurat) and UMAP generation were performed. All specific parameters and settings are documented in our code (https://github.com/betts-lab/asapseq-hiv-art).

### Initial ADT data processing

Sequencing reads were converted to fastq files and demultiplexed using bcl2fastq (Illumina). Reads were then inputted into kallisto bustools (Melsted et al., 2021) to generate cell barcode by feature matrices. These matrices were loaded into R and checked for empty droplets (i.e. background noise) using the emptyDrops function from the DropletUtils (Lun et al., 2019) package with a lower bound of 500 UMIs. Barcodes with false discovery rates < 0.01 were kept as genuine cells. This filtered matrix was then loaded into Seurat (v4.1.1) (Hao et al., 2021) for centered log ratio transformation with a scale factor of 10000. Harmony (Korsunsky et al., 2019) was used for batch correction if multiple 10X GEM wells were used for the same individual. A wilcoxon test (right-tailed) was used to select for features with a count distribution that was significantly different than background signal detected from isotype control antibodies, as analyzed similarly in a prior report (Swanson et al., 2021). A feature was not included in downstream analyses if p > 0.05 in at least 4 isotype controls.

### Identification of HIV+ cells

We built a custom Python pipeline (hiv-haystack; https://github.com/betts-lab/hiv-haystack) built and derived from epiVIA (Wang et al., 2020) to identify proviral reads from the BAM output file of cellranger-atac count. BAM records are parsed and selected into three groups: 1) both mates aligning to provirus, 2) both mates in host with a soft align clip, and 3) one mate in provirus and one mate in host. In scenario 1, the original alignments were kept as bona fide proviral alignments. If there was a soft clip present, we aligned it to the host genome using bwa mem as done in epiVIA to assess if an integration site is present. In scenario 2, any host soft clips were checked for exact matching to LTR sequences to allow for integration site detection. This was only allowed if a LTR alignment was reported (checked with BLAST against HXB2 5’ LTR). In scenario 3, the proviral mate was saved as a proviral fragment. If either mate had a soft clip, integration site checking is performed similar to scenarios 1 and 2. In all scenarios, only exact integration sites with exact nucleotide position are kept for downstream analyses. Our program also differs from epiVIA in that hiv-haystack allows for alignments to multiple sequences (from near full-length genomes and single genome sequencing of HIV Env) which enhances the detection of donor-derived sequences.

### Downstream data processing

After initial ATAC and ADT processing, barcodes that passed quality control metrics in both modalities were selected for downstream analyses. Clusters were annotated from a base panel of ADT expression patterns, imputed gene scores from ATAC, and differentially expressed markers present in any given cluster over all other cells. UMAP embeddings in lower dimension were generated using ArchR’s latent semantic indexing implementation. Manual annotations were then used to subset cells for downstream ADT differential expression analysis between HIV+ and HIV- cells using Seurat FindMarkers (using “DESeq2” for the findMarkerMethod parameter) to limit false discoveries (Squair et al., 2021) and the subset of features that were significantly different from background expression. For downstream ATAC differential expression analysis, manual clusters were grouped based on CD4+/- and HIV+/- (and by memory phenotype for the *in vitro* experiment) and pseudo-bulk replicates were generated. Clusters with less than 0.5% of the population were filtered out due to annotation difficulty. Peaks were called using the default 501-bp fixed-width method with ArchR and MACS2 (Zhang et al., 2008). Differential peaks were assessed using a wilcoxon test as implemented by ArchR’s getMarkerFeatures with bias parameters set to “c(TSSEnrichment”, “log10(nFrags”))” as recommended by ArchR. Motif annotations of differential peaks were added using addMotifAnnotations from the CISBP dataset. For experiments with lower numbers of HIV+ cells, we used ArchR’s implementation of chromVAR to determine differential motifs. getMarkerFeatures was called to determine differentially expressed motifs using the chromVAR deviation scores (i.e. z-score).

### Supervised machine learning

Samples were split into a 70% training and 30% testing dataset using the caTools package in R. Logistic regression models were developed using the glm function with parameters “family = binomial(link = ‘logit’)”. Naive Bayesian models were constructed using the naivebayes package in R. Random forest models were constructed using the randomForest package in R with ntree = 4000. Classifier performances (true positive rate and false positive rate) were calculated using the ROCR package in R and plotted using ggplot2.

### NFL SGS of proviral DNA from resting CD4+ T cells

DNA was extracted from multiples of 4 million resting CD4+ T cells according to the manufacturer’s instructions (QIAamp DNA Mini and Blood Mini Kit, QIAGEN). Amplification of NFL genomes was performed by limiting dilution, nested PCR using Platinum Taq High Fidelity Polymerase (Life Technologies, Thermo Fisher Scientific) as previously described (Bar et al., 2016), adapted for NFL genomes with primary forward and reverse primers, respectively 5′-AAATCTCTAGCAGTGGCGCCCGAACAG-3′ and 5′-TGAGGGATCTCTAGTTACCAGAGTC-3′ followed by nested forward and reverse primers, respectively 5′-GCGGAGGCTAGAAGGAGAGAGATGG-3′ and 5′-GCACTCAAGGCAAGCTTTATTGAGGCTTA-3′. Amplicons of appropriate size were directly sequenced on the Illumina MiSeq platform, inspected for evidence of priming from multiple templates, stop codons, large deletions, or introduction of PCR error in early cycles; a threshold of 85% identity at each nucleotide position was used.

### HIV custom sequence annotation

HIV custom sequences from untreated and treated PLWH were loaded into the Gene Cutter tool (HIV LANL Database) for alignment to HIV genes/regions. The output was then loaded into our custom tool, called geneCutterParser (https://github.com/wuv21/geneCutterParser), to extract the coordinate locations for any alignments. The coordinate locations were then used for alignment graphs in the supplemental figures.

### Graphics

All figures were made in R with the following packages: gridExtra, ggplot2 (Wickham,2016), ArchR, and patchwork. Graphical abstract cell icons were made with Biorender.com. All code to produce graphs can be found in the study GitHub repository.

## Author contributions

VHW, KJB, LAV, and MRB conceptualized experiments. PT, LJM, and KJB recruited and organized sample collection and cell preparation that were used for the study. VHW and JMLN performed the ASAPseq and flow experiments. JJ, FM, and KJB conducted and compiled the consensus viral sequencing. VHW and SN performed the ASAPseq data analysis. VHW, LAV, and MRB wrote the manuscript.

## Acknowledgements

We thank the individuals who donated samples to our study. We would also like to thank members of the Betts, Vella, and Bar labs for their assistance, Andrew Wells and James Pippin for assistance with NovaSeq sequencing, the University of Pennsylvania Next-Generation Sequencing Core, and the University of Pennsylvania Human Immunology Core. Additional thanks to Wenliang Wang, Golnaz Vahedi, and Brandon Keele for helpful discussions regarding proviral alignment and software. MRB is supported by the NIH grant (U19-A1-149680-02). Both MRB and KJB are supported by the NIH grant (P01-AI31338). Both MRB and LAV are further supported by the Penn Center for AIDS Research (P30-AI045008). LAV is supported by NIH grant K08-AI136660. VHW is supported by NIH grant (T32-AI007632). MRB, KJB, PT, and LJM are supported by UM-1AI164570 (BEAT-HIV Collaboratory) which is co-supported by the National Institute of Allergies and Infectious Diseases (NIAID), the National Institute of Mental Health (NIMH), the National Institute of Neurological Disorders and Stroke (NINDS), the National Institute on Drug Abuse (NIDA), and the Robert I. Jacobs Fund of The Philadelphia Foundation. LJM is also supported by the Herbert Kean, M.D., Family Professorship.

