## Supplemental Figures for "Profound phenotypic and epigenetic heterogeneity of the HIV-1 infected CD4+ T cell reservoir"

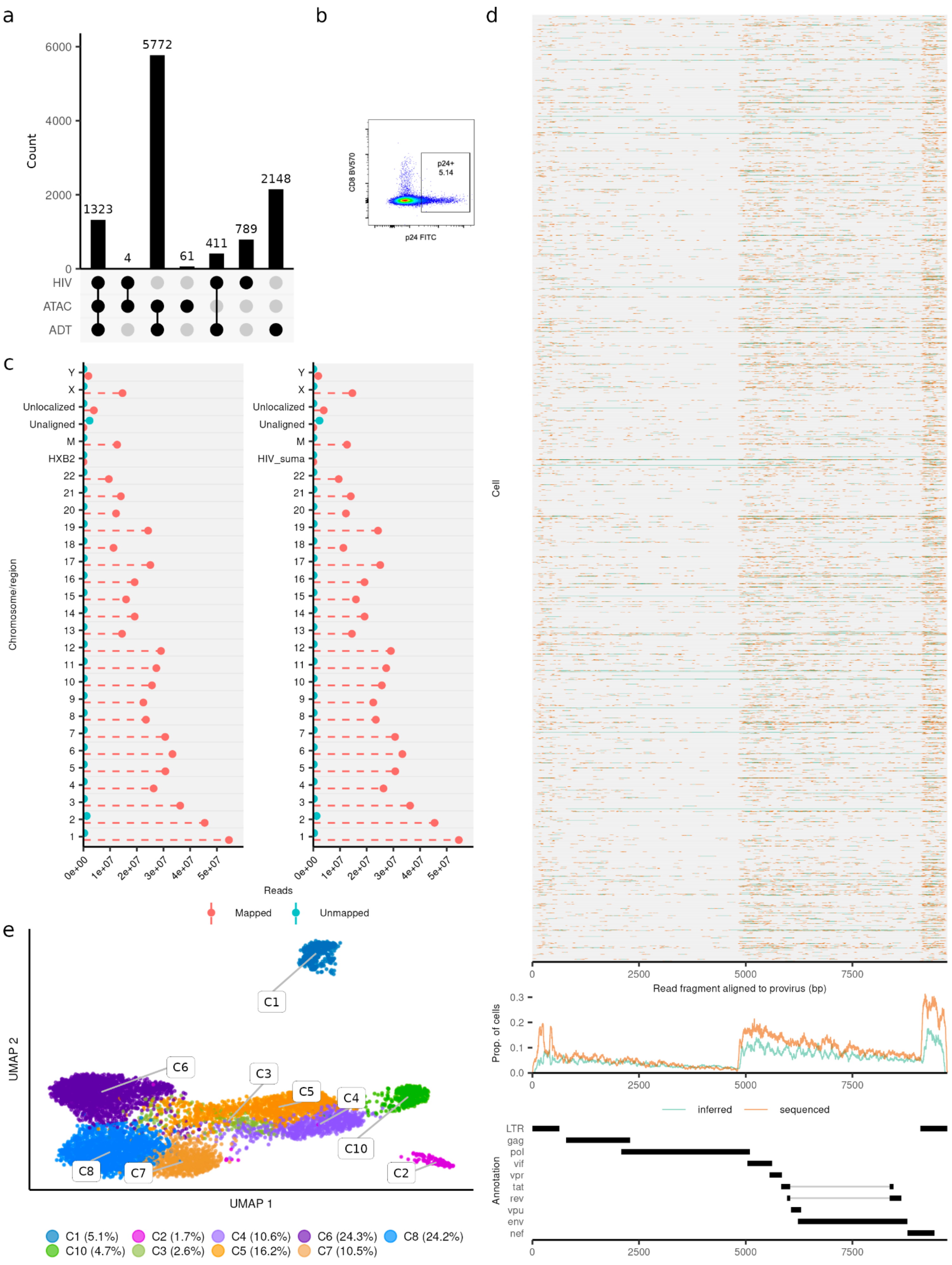

a
b
c
d
e
Count
1323
4
5772
61
411
789
2148
HIV
ATAC
ADT
CD8 BV570
p24 FITC
p24+
5.14
Chromosome/region
Unlocalized
Unaligned
HXB2
HIV_suma
Reads
Mapped
Unmapped
Cell
Read fragment aligned to provirus (bp)
Prop. of cells
inferred
sequenced
Annotation
LTR
gag
pol
vif
vpr
tat
rev
vpu
env
nef
UMAP 1
UMAP 2
C1 (5.1%)
C2 (1.7%)
C4 (10.6%)
C6 (24.3%)
C8 (24.2%)
C10 (4.7%)
C3 (2.6%)
C5 (16.2%)
C7 (10.5%)

**Supplemental Figure 1: Properties of ASAPseq library for *in vitro* model.** (A) UpSet plot of unique cell barcodes that were collected from each modality (ATAC versus ADT) and whether or not the barcode was associated with proviral reads (HIV). Barcodes that passed ATAC and ADT quality checks (see Methods) were used for downstream analyses. (B) Flow cytometry plot of cell culture before conducting ASAPseq analysis. X-axis is p24 while Y-axis is CD8. Value in the highlighted box is the percent of total live singlets that are p24+. (C) Reported mapped and unmapped read-segments by chromosome (as determined by samtools idxstats) from alignment of ASAPseq dataset of uninfected PBMC (Mimitou et al., 2021) to chimeric reference genomes with HXB2 (left) or SUMA (right). HIV genomes were added as a separate chromosome during creation of the chimeric reference genome. (D) (top) Sequenced regions that are aligned by bwa mem to the proviral genome (SUMA) and recovered by hiv-haystack. Each row is a cell and each column is a base pair spanning the proviral genome. Regions in orange indicate actual reported coverage while regions in blue indicate inferred coverage if provirus was intact as paired-end sequencing can only obtain at most 50bp from either end of the genomic/transposed fragment if the genomic fragment is > 50bp. Many LTR alignments can be ambiguous and it is unclear whether the actual read is in the 3' LTR or 5' LTR. The primary alignment from bwa-mem is recorded here. (middle) Proportion of coverage is reported across all cells spanning the entire proviral genome. (bottom) Genome map of SUMA. (E) UMAP representation of the ATAC component with numeric labeling prior to manual annotation.

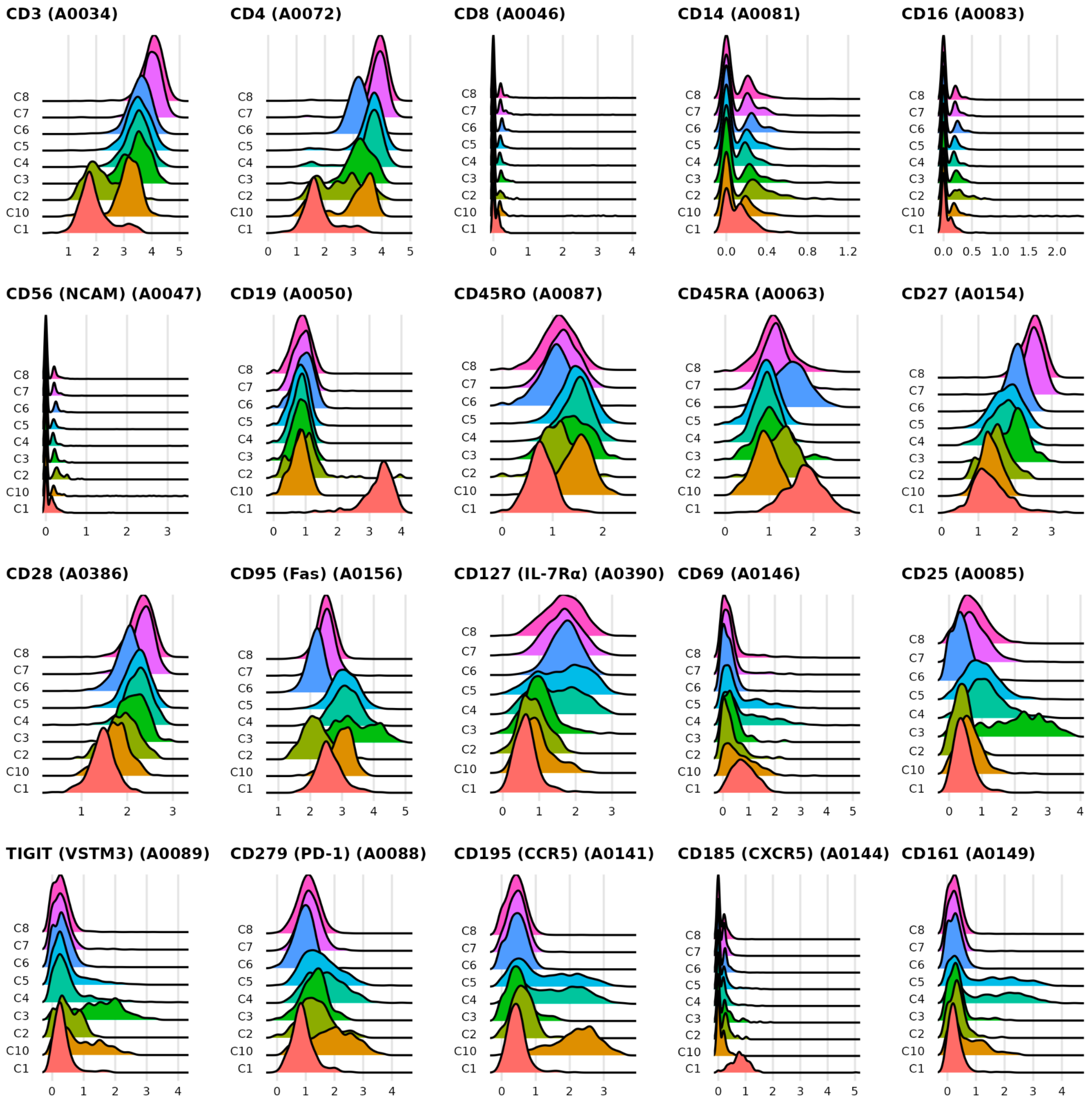


**Supplemental Figure 2: ADT base panel for *in vitro* model.** Each subplot shows the ADT signal for a specific surface antigen for each cluster as seen in Supplemental Figure 1E. X-axis values are normalized count values as processed via Seurat.

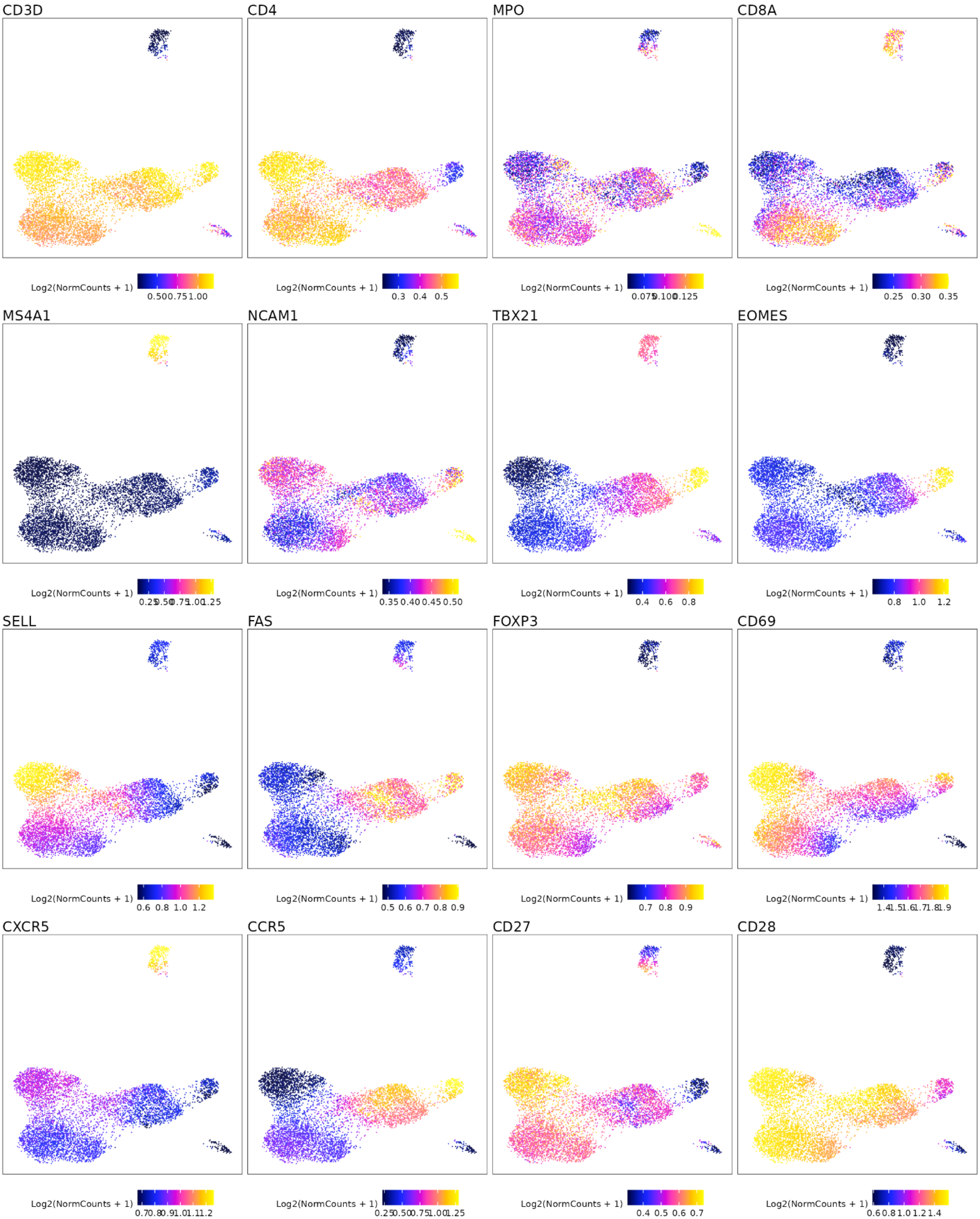
CD3D
CD4
MPO
CD8A
MS4A1
NCAM1
TBX21
EOMES
SELL
FAS
FOXP3
CD69
CXCR5
CCR5
CD27
CD28
Log2(NormCounts + 1)
0.3 0.4 0.5
0.25 0.30 0.35
0.35 0.40 0.45 0.50
0.4 0.6 0.8
0.8 1.0 1.2
0.6 0.8 1.0 1.2
0.5 0.6 0.7 0.8 0.9
0.7 0.8 0.9
0.4 0.5 0.6 0.7

**Supplemental Figure 3: ATAC base panel for *in vitro* model.** Each subplot shows the imputed gene activity score overlaid on the UMAP coordinate space as seen in Supplementary Figure 1E. Gene activity score was calculated by ArchR and imputed using MAGIC to aid in visual interpretation as recommended by ArchR.

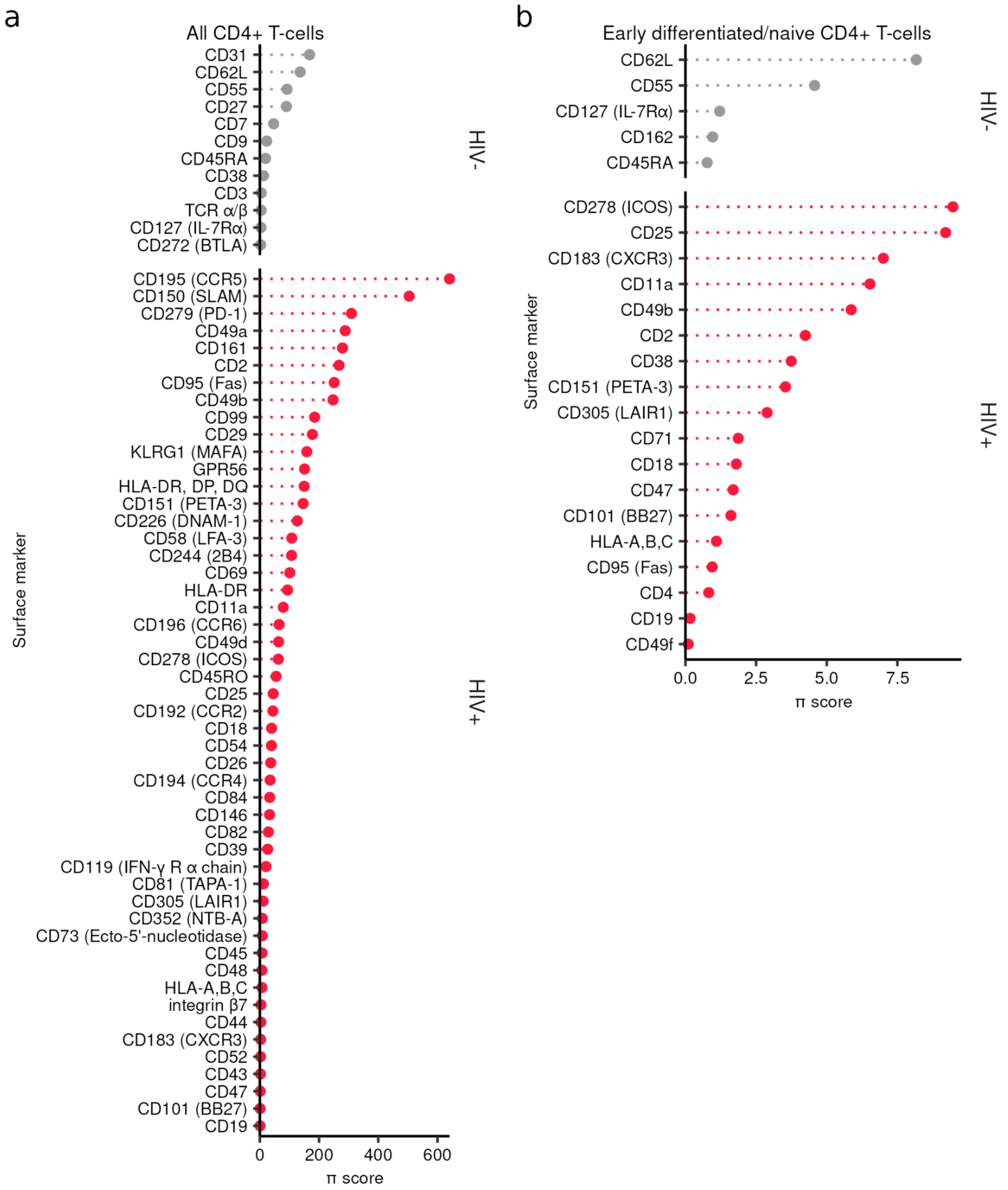


**Supplementary Figure 4: Differential expression of select antigens for *in vitro* model.** Significant surface markers (adjusted p value < 0.05) enriched in HIV- or HIV+ cells are shown for (A) all CD4+ T-cells and (B) early differentiated CD4+ T-cells. Gray indicates markers for HIV- cells while gray indicates markers for HIV+ cells. π-score was calculated as log10(adjusted p value) * log2(fold change).

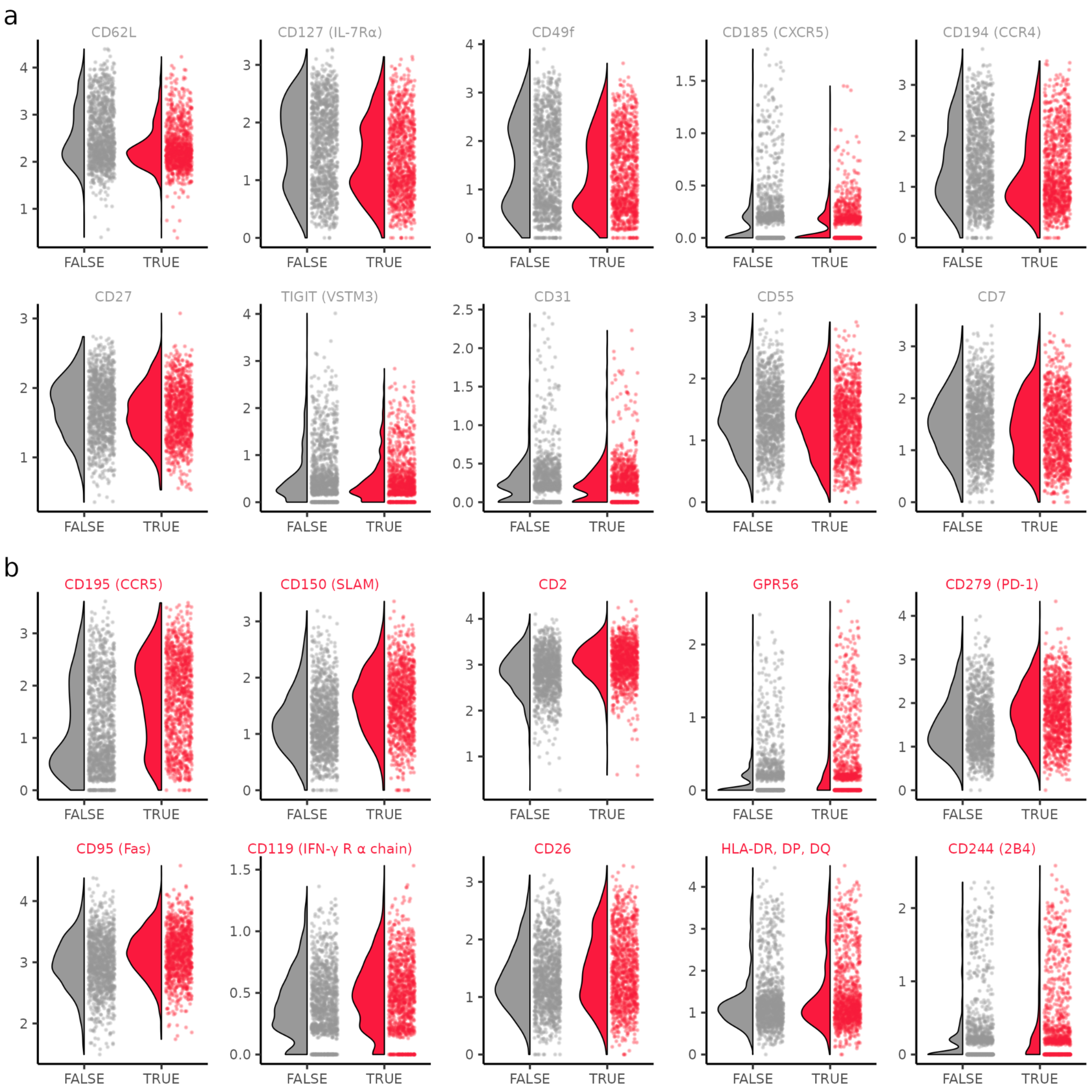


**Supplementary Figure 5: Differential expression of select antigens in activated T-cells for *in vitro* model.** Violin-scatter plots are shown for the top 10 surface markers from Figure 1E that are enriched in (A) activated HIV- cells and (B) activated HIV+ cells. Grey color (FALSE) indicates HIV- cells while red color (TRUE) indicates HIV+ cells. Markers are ordered from left to right; top to bottom by order of decreasing |π-score| as seen in Figure 1E.

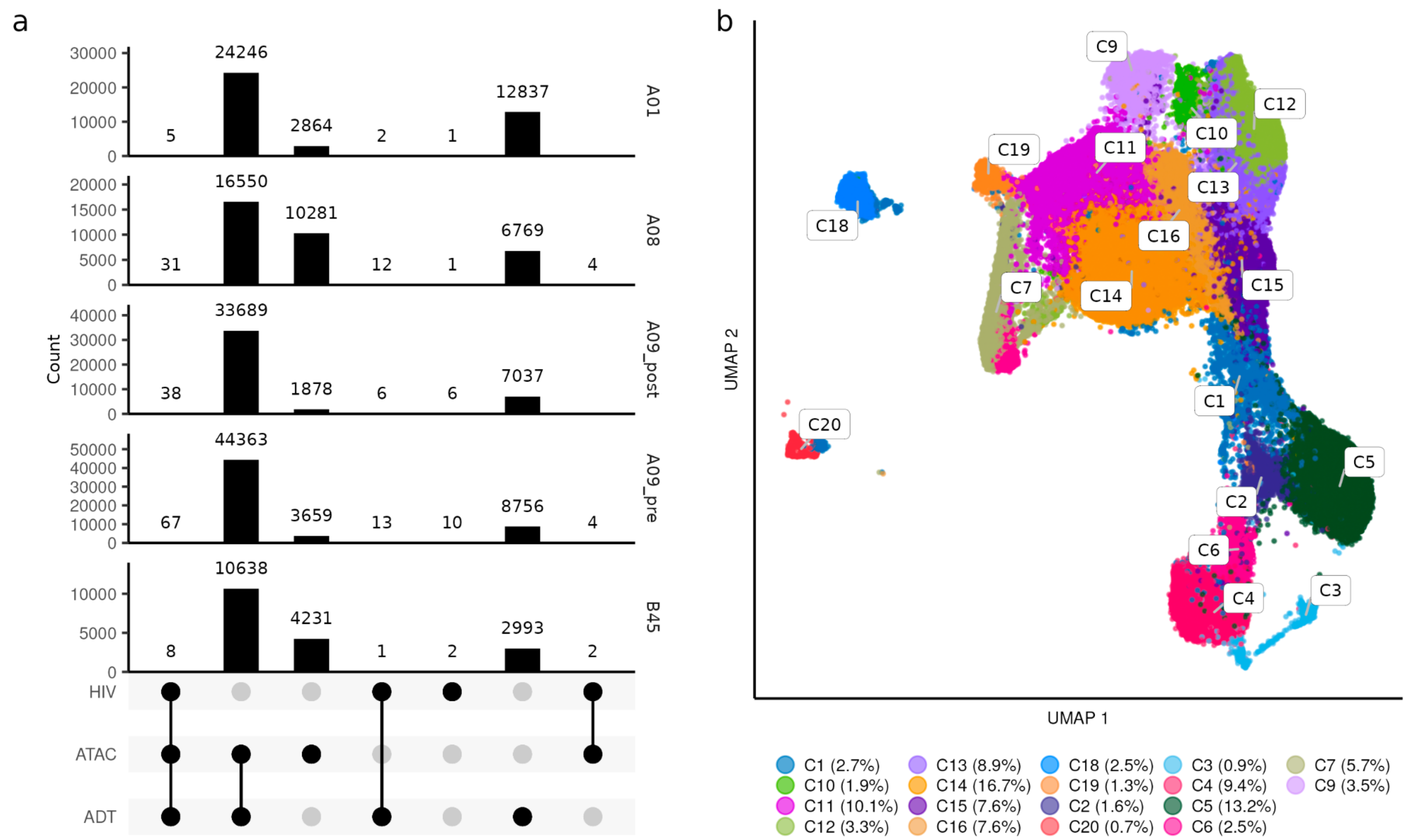


**Supplementary Figure 6: Properties of ASAPseq library during treated infection.** (A) UpSet plot of unique cell barcodes that were collected from each modality (ATAC versus ADT) and detection of proviral reads (HIV) separated by individual. (B) UMAP representation of the ATAC component with numeric labeling prior to manual annotation.

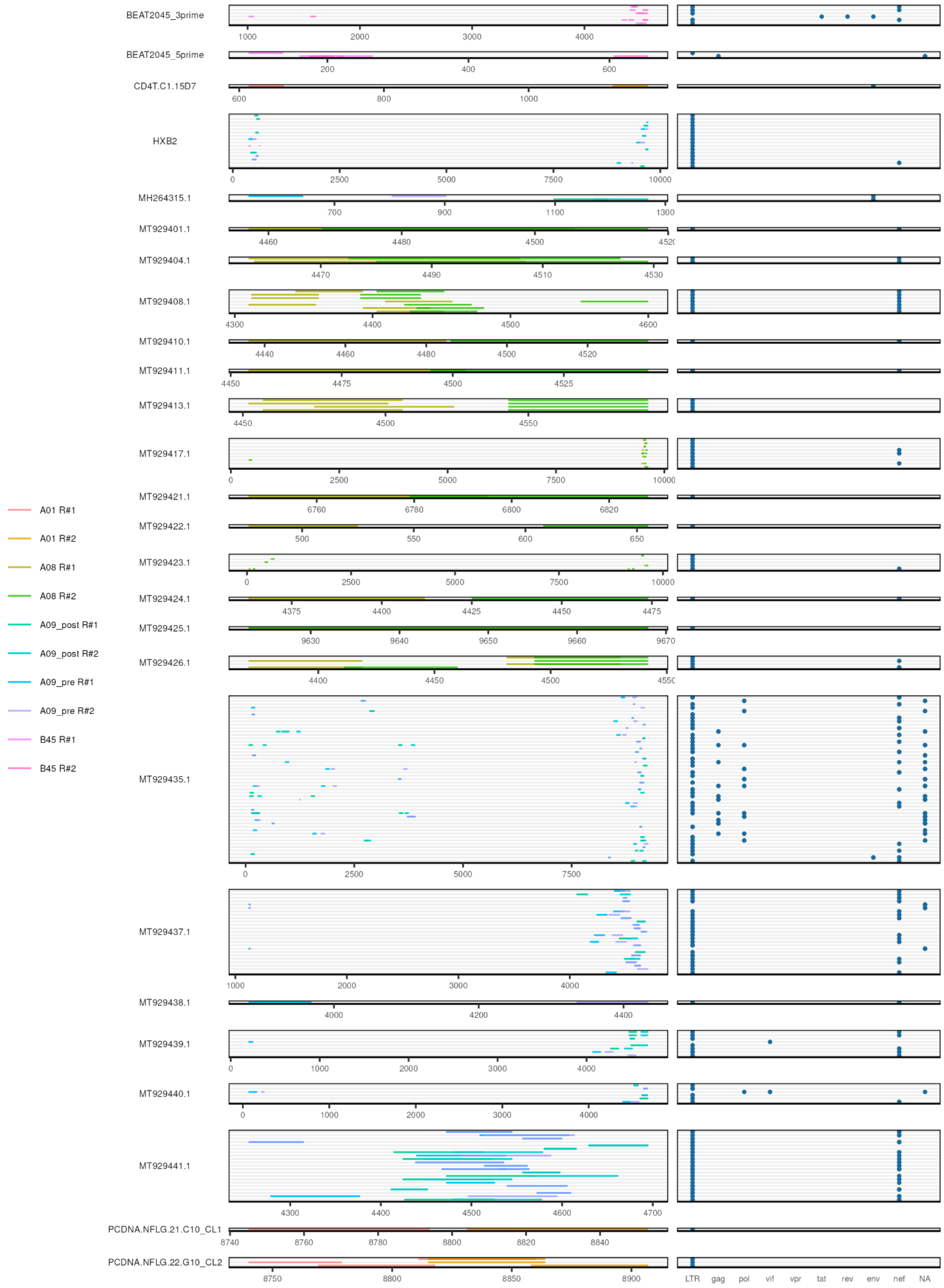

BEAT2045_3prime
BEAT2045_5prime
CD4T.C1.15D7
HXB2
MH264315.1
MT929401.1
MT929404.1
MT929408.1
MT929410.1
MT929411.1
MT929413.1
MT929417.1
MT929421.1
MT929422.1
MT929423.1
MT929424.1
MT929425.1
MT929426.1
MT929435.1
MT929437.1
MT929438.1
MT929439.1
MT929440.1
MT929441.1
PCDNA.NFLG.21.C10_CL1
PCDNA.NFLG.22.G10_CL2
A01 R#1
A01 R#2
A08 R#1
A08 R#2
A09_post R#1
A09_post R#2
A09_pre R#1
A09_pre R#2
B45 R#1
B45 R#2
LTR gag pol vif vpr tat rev env nef NA

**Supplementary Figure 7: HIV alignment for ART-PLWH samples.** (left) Sequenced regions that are aligned by bwa mem to the proviral genome (autologous + HXB2) and recovered by hiv-haystack. Each row is a cell and each column is a base pair spanning different autologous sequences. Tracks are separated by autologous sequence (sequence name is shown on the left in gray box). (right) For each row, the coverage is labeled by its annotation (as determined from Gene Cutter).

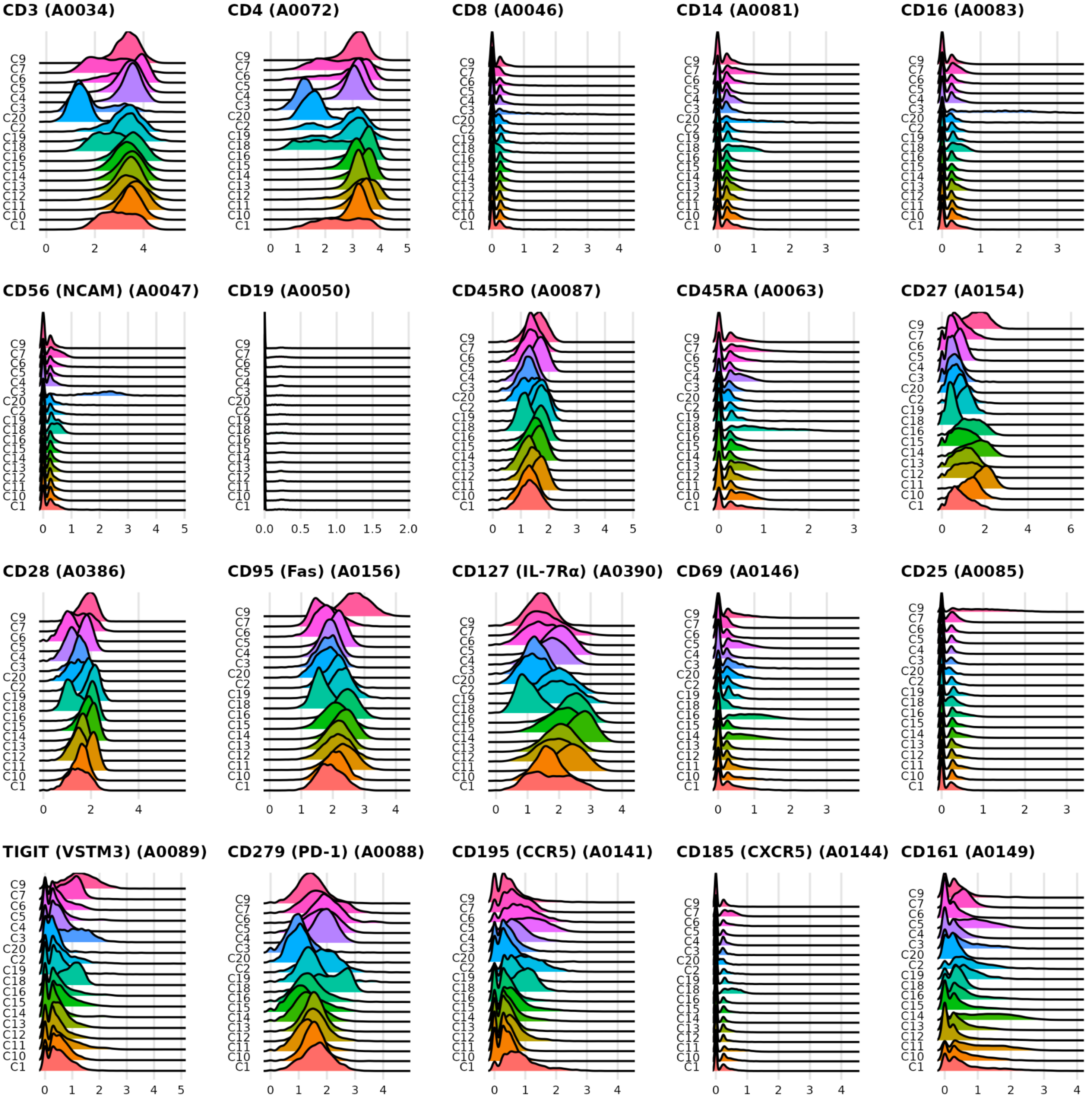


**Supplementary Figure 8: ADT base panel during treated infection.** Each subplot shows the ADT signal for a specific surface antigen for each cluster as seen in Supplemental Figure 6B. X-axis values are normalized count values as processed via Seurat.

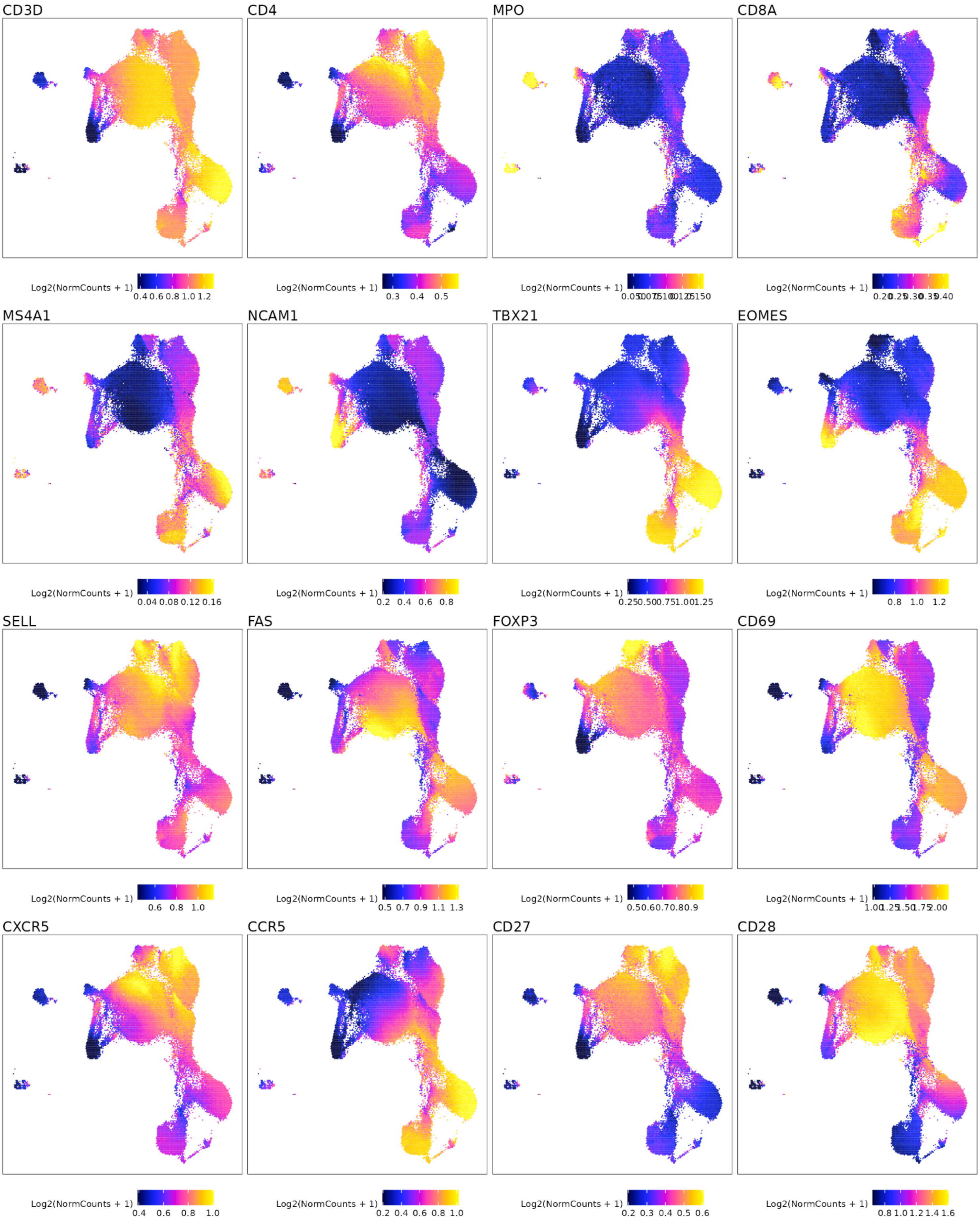

CD3D
CD4
MPO
CD8A
Log2(NormCounts + 1)
0.4 0.6 0.8 1.0 1.2
0.3 0.4 0.5
MS4A1
NCAM1
TBX21
EOMES
0.04 0.08 0.12 0.16
0.2 0.4 0.6 0.8
0.8 1.0 1.2
SELL
FAS
FOXP3
CD69
0.6 0.8 1.0
0.5 0.7 0.9 1.1 1.3
CXCR5
CCR5
CD27
CD28
0.4 0.6 0.8 1.0
0.2 0.4 0.6 0.8 1.0
0.2 0.3 0.4 0.5 0.6
0.8 1.0 1.2 1.4 1.6

**Supplementary Figure 9: ATAC base panel during treated infection.** Each subplot shows the imputed gene activity score overlaid on the UMAP coordinate space as seen in Supplementary Figure 6B. Gene activity score was calculated by ArchR and imputed using MAGIC to aid in visual interpretation as recommended by ArchR.

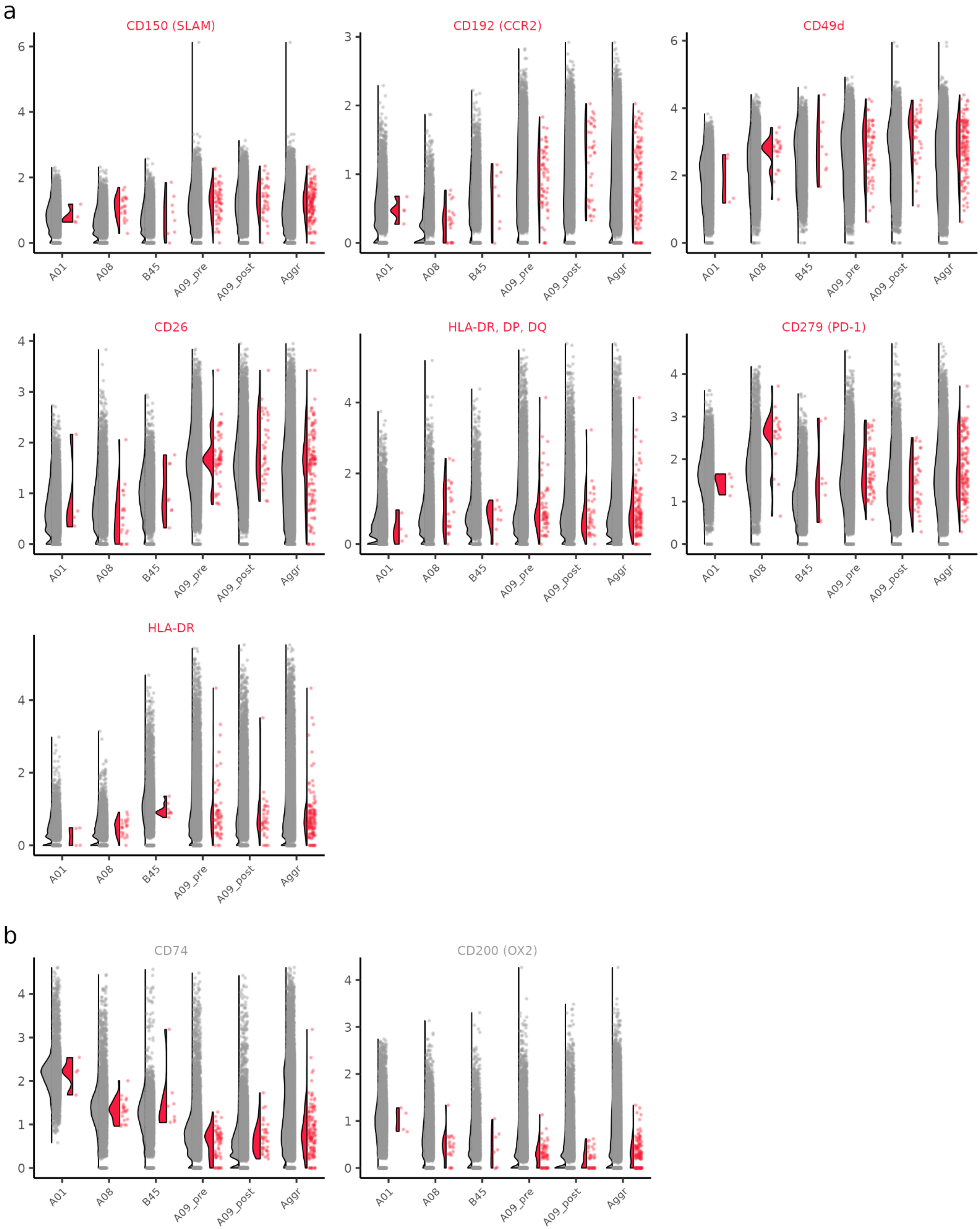

a
CD150 (SLAM)
CD192 (CCR2)
CD49d
CD26
HLA-DR, DP, DQ
CD279 (PD-1)
HLA-DR
b
CD74
CD200 (OX2)
A01
A08
B45
A09_pre
A09_post
Aggr

**Supplementary Figure 10: Differential expression of select antigens during treated infection.** Violin-scatter plots are shown for all significantly expressed (adjusted p value < 0.05; Wilcoxon) surface markers from Figure 4B that are enriched in (A) all CD4+ HIV+ T-cells and (B) all CD4+ HIV- T-cells. Grey color indicates HIV- cells while red color indicates HIV+ cells. Markers are ordered from left to right; top to bottom by order of decreasing |π-score| as seen in Figure 4B. Cells are separated by individual and the aggregate data (across individuals) is also shown.
