## Supplemental Tables for "Profound phenotypic and epigenetic heterogeneity of the HIV-1 infected CD4+ T cell reservoir"

**Supplemental Table 1:** Catalog of TotalSeqA antibodies used for study.

| DNA_ID | Description | Clone | Barcode |
|---|---|---|---|
| A0006 | CD86 | IT2.2 | GTCTTTGTCAGTGCA |
| A0007 | CD274 (B7-H1, PD-L1) | 29E.2A3 | GTTGTCCGACAATAC |
| A0020 | CD270 (HVEM, TR2) | 122 | TGATAGAAACAGACC |
| A0023 | CD155 (PVR) | SKII.4 | ATCACATCGTTGCCA |
| A0024 | CD112 (Nectin-2) | TX31 | AACCTTCCGTCTAAG |
| A0026 | CD47 | CC2C6 | GCATTCTGTCACCTA |
| A0029 | CD48 | BJ40 | CTACGACGTAGAAGA |
| A0031 | CD40 | 5C3 | CTCAGATGGAGTATG |
| A0032 | CD154 | 24-31 | GCTAGATAGATGCAA |
| A0033 | CD52 | HI186 | CTTTGTACGAGCAAA |
| A0034 | CD3 | UCHT1 | CTCATTGTAACTCCT |
| A0046 | CD8 | SK1 | GCGCAACTTGATGAT |
| A0047 | CD56 (NCAM) | 5.1H11 | TCCTTTCCTGATAGG |
| A0050 | CD19 | HIB19 | CTGGGCAATTACTCG |
| A0052 | CD33 | P67.6 | TAACTCAGGGCCTAT |
| A0053 | CD11c | S-HCL-3 | TACGCCTATAACTTG |
| A0058 | HLA-A,B,C | W6/32 | TATGCGAGGCTTATC |
| A0063 | CD45RA | HI100 | TCAATCCTTCCGCTT |
| A0064 | CD123 | 6H6 | CTTCACTCTGTCAGG |
| A0066 | CD7 | CD7-6B7 | TGGATTCCCGGACTT |
| A0070 | CD49f | GoH3 | TTCCGAGGATGATCT |
| A0071 | CD194 (CCR4) | L291H4 | AGCTTACCTGCACGA |
| A0072 | CD4 | RPA-T4 | TGTTCCCGCTCAACT |
| A0073 | CD44 | IM7 | TGGCTTCAGGTCCTA |
| A0081 | CD14 | M5E2 | TCTCAGACCTCCGTA |
| A0083 | CD16 | 3G8 | AAGTTCACTCTTTGC |
| A0085 | CD25 | BC96 | TTTGTCCTGTACGCC |
| A0087 | CD45RO | UCHL1 | CTCCGAATCATGTTG |
| A0088 | CD279 (PD-1) | EH12.2H7 | ACAGCGCCGTATTTA |
| A0089 | TIGIT (VSTM3) | A15153G | TTGCTTACCGCCAGA |
| A0090 | Mouse IgG1, κ isotype Ctrl | MOPC-21 | GCCGGACGACATTAA |
| A0091 | Mouse IgG2a, κ isotype Ctrl | MOPC-173 | CTCCTACCTAAACTG |
| A0092 | Mouse IgG2b, κ isotype Ctrl | MPC-11 | ATATGTATCACGCGA |
| A0095 | Rat IgG2b, κ Isotype Ctrl | RTK4530 | GATTCTTGACGACCT |
| A0100 | CD20 | 2H7 | TTCTGGGTCCCTAGA |
| A0101 | CD335 (NKp46) | 9E2 | ACAATTTGAACAGCG |
| A0124 | CD31 | WM59 | ACCTTTATGCCACGG |
| A0127 | Podoplanin | NC-08 | GGTTACTCGTTGTGT |
| A0134 | CD146 | P1H12 | CCTTGGATAACATCA |
| A0136 | IgM | MHM-88 | TAGCGAGCCCGTATA |
| A0138 | CD5 | UCHT2 | CATTAACGGGATGCC |
| A0140 | CD183 (CXCR3) | G025H7 | GCGATGGTAGATTAT |
| A0141 | CD195 (CCR5) | J418F1 | CCAAAGTAAGAGCCA |
| A0142 | CD32 | FUN-2 | GCTTCCGAATTACCG |
| A0143 | CD196 (CCR6) | G034E3 | GATCCCTTTGTCACT |
| A0144 | CD185 (CXCR5) | J252D4 | AATTCAACCGTCGCC |
| A0145 | CD103 (Integrin αE) | Ber-ACT8 | GACCTCATTGTGAAT |
| A0146 | CD69 | FN50 | GTCTCTTGGCTTAAA |
| A0147 | CD62L | DREG-56 | GTCCCTGCAACTTGA |
| A0149 | CD161 | HP-3G10 | GTACGCAGTCCTTCT |
| A0151 | CD152 (CTLA-4) | BNI3 | ATGGTTCACGTAATC |
| A0152 | CD223 (LAG-3) | 11C3C65 | CATTTGTCTGCCGGT |

*(continued)*

| DNA_ID | Description | Clone | Barcode |
|---|---|---|---|
| A0153 | KLRG1 (MAFA) | SA231A2 | CTTATTTCCTGCCCT |
| A0154 | CD27 | O323 | GCACTCCTGCATGTA |
| A0155 | CD107a (LAMP-1) | H4A3 | CAGCCCACTGCAATA |
| A0156 | CD95 (Fas) | DX2 | CCAGCTCATTAGAGC |
| A0158 | CD134 (OX40) | Ber-ACT35 (ACT35) | AACCCACCGTTGTTA |
| A0159 | HLA-DR | L243 | AATAGCGAGCAAGTA |
| A0160 | CD1c | L161 | GAGCTACTTCACTCG |
| A0161 | CD11b | ICRF44 | GACAAGTGATCTGCA |
| A0162 | CD64 | 10.1 | AAGTATGCCCTACGA |
| A0163 | CD141 (Thrombomodulin) | M80 | GGATAACCGCGCTTT |
| A0165 | CD314 (NKG2D) | 1D11 | CGTGTTTGTTCCTCA |
| A0167 | CD35 | E11 | ACTTCCGTCGATCTT |
| A0168 | CD57 | QA17A04 | AACTCCCTATGGAGG |
| A0170 | CD272 (BTLA) | MIH26 | GTTATTGGACTAAGG |
| A0171 | CD278 (ICOS) | C398.4A | CGCGCACCCATTAAA |
| A0172 | CD275 (B7-H2, B7-RP1, ICOSL) | 9F.8A4 | GTTAGTGTTAGCTTG |
| A0174 | CD58 (LFA-3) | TS2/9 | GTTCCTATGGACGAC |
| A0176 | CD39 | A1 | TTACCTGGTATCCGT |
| A0179 | CX3CR1 | K0124E1 | AGTATCGTCTCTGGG |
| A0180 | CD24 | ML5 | AGATTCCTTCGTGTT |
| A0181 | CD21 | Bu32 | AACCTAGTAGTTCGG |
| A0185 | CD11a | TS2/4 | TATATCCTTGTGAGC |
| A0187 | CD79b (Igβ) | CB3-1 | ATTCTTCAACCGAAG |
| A0189 | CD244 (2B4) | C1.7 | TCGCTTGGATGGTAG |
| A0206 | CD169 (Sialoadhesin, Siglec-1) | 7-239 | TACTCAGCGTGTTTG |
| A0214 | integrin β7 | FIB504 | TCCTTGGATGTACCG |
| A0215 | CD268 (BAFF-R) | 11C1 | CGAAGTCGATCCGTA |
| A0216 | CD42b | HIP1 | TCCTAGTACCGAAGT |
| A0217 | CD54 | HA58 | CTGATAGACTTGAGT |
| A0218 | CD62P (P-Selectin) | AK4 | CCTTCCGTATCCCTT |
| A0219 | CD119 (IFN-γ R α chain) | GIR-208 | TGTGTATTCCCTTGT |
| A0224 | TCR α/β | IP26 | CGTAACGTAGAGCGA |
| A0236 | Rat IgG1, κ isotype Ctrl | RTK2071 | ATCAGATGCCCTCAT |
| A0237 | Rat IgG1, λ Isotype Ctrl | G0114F7 | GGGAGCGATTCAACT |
| A0238 | Rat IgG2a, κ Isotype Ctrl | RTK2758 | AAGTCAGGTTCGTTT |
| A0240 | Rat IgG2c, κ Isotype Ctrl | RTK4174 | TCCAGGCTAGTCATT |
| A0241 | Armenian Hamster IgG Isotype Ctrl | HTK888 | CCTGTCATTAAGACT |
| A0242 | CD192 (CCR2) | K036C2 | GAGTTCCCTTACCTG |
| A0246 | CD122 (IL-2Rβ) | TU27 | TCATTTCCTCCGATT |
| A0247 | CD267 (TACI) | 1A1 | AGTGATGGAGCGAAC |
| A0352 | FcεRIα | AER-37 (CRA-1) | CTCGTTTCCGTATCG |
| A0353 | CD41 | HIP8 | ACGTTGTGGCCTTGT |
| A0355 | CD137 (4-1BB) | 4B4-1 | CAGTAAGTTCGGGAC |
| A0357 | CD43 | CD43-10G7 | GATTAACCAGCTCAT |
| A0358 | CD163 | GHI/61 | GCTTCTCCTTCCTTA |
| A0359 | CD83 | HB15e | CCACTCATTTCCGGT |
| A0364 | CD13 | WM15 | TTTCAACGCCCTTTC |
| A0367 | CD2 | TS1/8 | TACGATTTGTCAGGG |
| A0368 | CD226 (DNAM-1) | 11A8 | TCTCAGTGTTTGTGG |
| A0369 | CD29 | TS2/16 | GTATTCCCTCAGTCA |
| A0370 | CD303 (BDCA-2) | 201A | GAGATGTCCGAATTT |
| A0371 | CD49b | P1E6-C5 | GCTTTCTTCAGTATG |
| A0372 | CD61 | VI-PL2 | AGGTTGGAGTAGACT |

(continued)

| DNA_ID | Description | Clone | Barcode |
|---|---|---|---|
| A0373 | CD81 (TAPA-1) | 5A6 | GTATCCTTCCTTGGC |
| A0383 | CD55 | JS11 | GCTCATTACCCATTA |
| A0384 | IgD | IA6-2 | CAGTCTCCGTAGAGT |
| A0385 | CD18 | TS1/18 | TATTGGGACACTTCT |
| A0386 | CD28 | CD28.2 | TGAGAACGACCCTAA |
| A0389 | CD38 | HIT2 | TGTACCCGCTTGTGA |
| A0390 | CD127 (IL-7Rα) | A019D5 | GTGTGTTGTCCTATG |
| A0391 | CD45 | HI30 | TGCAATTACCCGGAT |
| A0393 | CD22 | S-HCL-1 | GGGTTGTTGTCTTTG |
| A0394 | CD71 | CY1G4 | CCGTGTTCCTCATTA |
| A0396 | CD26 | BA5b | GGTGGCTAGATAATG |
| A0398 | CD115 (CSF-1R) | 9-4D2-1E4 | AATCACGGTCCTTGT |
| A0404 | CD63 | H5C6 | GAGATGTCTGCAACT |
| A0406 | CD304 (Neuropilin-1) | 12C2 | GGACTAAGTTTCGTT |
| A0407 | CD36 | 5-271 | TTCTTTGCCTTGCCA |
| A0408 | CD172a (SIRPα) | 15-414 | CGTGTTTAACTTGAG |
| A0419 | CD72 | 3F3 | CAGTCGTGGTAGATA |
| A0420 | CD158 (KIR2DL1/S1/S3/S5) | HP-MA4 | TATCAACCAACGCTT |
| A0446 | CD93 | VIMD2 | GCGCTACTTCCTTGA |
| A0447 | CD200 (OX2) | OX-104 | CACGTAGACCTTTGC |
| A0575 | CD49a | TS2/7 | ACTGATGGACTCAGA |
| A0576 | CD49d | 9F10 | CCATTCAACTTCCGG |
| A0577 | CD73 (Ecto-5’-nucleotidase) | AD2 | CAGTTCCTCAGTTCG |
| A0579 | CD9 | HI9a | GAGTCACCAATCTGC |
| A0581 | TCR Vα7.2 | 3C10 | TACGAGCAGTATTCA |
| A0582 | TCR Vδ2 | B6 | TCAGTCAGATGGTAT |
| A0586 | CD354 (TREM-1) | TREM-26 | TAGCCGTTTCCTTTG |
| A0590 | CD305 (LAIR1) | NKTA255 | ATTTCCATTCCCTGT |
| A0591 | LOX-1 | 15C4 | ACCCTTTACCGAATA |
| A0599 | CD158e1 (KIR3DL1, NKB1) | DX9 | GGACGCTTTCCTTGA |
| A0817 | CD109 | W7C5 | CACTTAACTCTGGGT |
| A0822 | CD142 | NY2 | CACTGCCGTCGATTA |
| A0830 | CD319 (CRACC) | 162.1 | AGTATGCCATGTCTT |
| A0845 | CD99 | 3B2/TA8 | ACCCGTCCCTAAGAA |
| A0853 | CLEC12A | 50C1 | CATTAGAGTCTGCCA |
| A0861 | CD151 (PETA-3) | 50-6 | CTTACCTAGTCATTC |
| A0864 | CD352 (NTB-A) | NT-7 | AGTTTCCACTCAGGC |
| A0866 | CLEC1B (CLEC2) | AYP1 | TGCCAGTATCACGTA |
| A0867 | CD94 | DX22 | CTTTCCGGTCCTACA |
| A0868 | IgE | MHE-18 | GGATGTACCGCGTAT |
| A0870 | CD150 (SLAM) | A12 (7D4) | GTCATTGTATGTCTG |
| A0871 | CD162 | KPL-1 | ATATGTCAGAGCACC |
| A0872 | CD84 | CD84.1.21 | CTCCCTAGTTCCTTT |
| A0894 | Ig light chain κ | MHK-49 | AGCTCAGCCAGTATG |
| A0896 | CD85j (ILT2) | GHI/75 | CCTTGTGAGGCTATG |
| A0897 | CD23 | EBVCS-5 | TCTGTATAACCGTCT |
| A0898 | Ig light chain λ | MHL-38 | CAGCCAGTAAGTCAC |
| A0902 | CD328 (Siglec-7) | 6-434 | CTTAGCATTTCACTG |
| A0912 | GPR56 | CG4 | GCCTAGTTTCCGTTT |
| A0920 | CD82 | ASL-24 | TCCCACTTCCGCTTT |
| A0923 | NKp80 | 5D12 | TATAGTTCCTCTGTG |
| A0931 | CD131 | 1C1 | CTGCATGAGACCAAA |

*(continued)*

| DNA_ID | Description | Clone | Barcode |
|---|---|---|---|
| A0935 | CD74 | LN2 | CTGTAGCATTTCCCT |
| A0940 | CD116 | 4H1 | ATGGACAGTTCGTGT |
| A0941 | CD37 | M-B371 | ACAGTCACTGGGCAA |
| A0944 | CD101 (BB27) | BB27 | CTACTTCCCTGTCAA |
| A1018 | HLA-DR, DP, DQ | Tü39 | AGCTACGAGCAGTAG |
| A1046 | CD88 (C5aR) | S5/1 | GCCGCATGAGAAACA |

**Supplemental Table 2:** Significant surface markers enriched in each cluster (over all other cells) for *in vitro* model.

| Cluster | Marker | log2(fold change) | Adjusted p value | π-score |
|---|---|---|---|---|
| C1 | CD22 | 1.40 | 2.0e-215 | 301.34 |
| C1 | CD72 | 1.52 | 1.0e-198 | 300.64 |
| C1 | CD20 | 1.26 | 2.5e-218 | 273.94 |
| C1 | CD40 | 1.17 | 6.4e-226 | 263.86 |
| C1 | HLA-DR | 1.20 | 2.2e-217 | 259.29 |
| C1 | CD19 | 1.18 | 5.6e-217 | 255.89 |
| C1 | CD32 | 1.14 | 5.3e-215 | 245.08 |
| C1 | HLA-DR, DP, DQ | 1.16 | 4.1e-212 | 244.45 |
| C1 | CD21 | 0.95 | 1.3e-240 | 227.59 |
| C1 | CD86 | 1.01 | 9.1e-209 | 210.48 |
| C1 | CD71 | 0.96 | 1.0e-204 | 196.72 |
| C1 | CD35 | 1.06 | 3.0e-155 | 164.48 |
| C1 | IgD | 1.26 | 3.2e-128 | 161.24 |
| C1 | CD85j (ILT2) | 0.78 | 2.2e-197 | 153.44 |
| C1 | CD39 | 0.75 | 1.9e-201 | 150.80 |
| C1 | CD185 (CXCR5) | 0.66 | 1.7e-211 | 138.28 |
| C1 | CD63 | 0.64 | 1.2e-167 | 106.64 |
| C1 | CD150 (SLAM) | 0.71 | 4.8e-145 | 103.18 |
| C1 | CD54 | 0.59 | 4.7e-174 | 101.59 |
| C1 | CD267 (TACI) | 0.62 | 1.0e-151 | 93.58 |
| C1 | CD79b (Igβ) | 0.57 | 7.7e-140 | 79.30 |
| C1 | CD82 | 0.36 | 2.2e-160 | 57.26 |
| C1 | CD119 (IFN-γ R α chain) | 0.47 | 7.4e-116 | 53.72 |
| C1 | CD74 | 0.35 | 9.3e-144 | 50.56 |
| C1 | CD45RA | 0.38 | 4.6e-114 | 42.61 |
| C1 | CD196 (CCR6) | 0.43 | 4.0e-92 | 38.88 |
| C1 | CD123 | 0.43 | 1.2e-88 | 37.49 |
| C1 | CD73 (Ecto-5'-nucleotidase) | 0.46 | 1.5e-80 | 36.40 |
| C1 | CD37 | 0.30 | 5.2e-102 | 30.09 |
| C1 | CD58 (LFA-3) | 0.32 | 2.0e-79 | 25.23 |
| C1 | CD1c | 0.29 | 2.2e-82 | 24.08 |
| C1 | CD69 | 0.34 | 3.5e-69 | 23.37 |
| C1 | CD319 (CRACC) | 0.31 | 2.7e-71 | 21.63 |
| C1 | IgM | 0.28 | 5.0e-62 | 17.24 |
| C1 | CD183 (CXCR3) | 0.29 | 2.4e-56 | 16.31 |
| C1 | CD272 (BTLA) | 0.26 | 6.5e-51 | 12.80 |
| C1 | CD83 | 0.29 | 6.2e-37 | 10.57 |
| C10 | CD244 (2B4) | 1.05 | 4.6e-218 | 227.73 |
| C10 | KLRG1 (MAFA) | 1.11 | 6.4e-173 | 190.52 |
| C10 | CD57 | 1.23 | 7.3e-134 | 163.63 |
| C10 | GPR56 | 0.84 | 4.6e-170 | 142.86 |
| C10 | CD195 (CCR5) | 0.90 | 3.1e-145 | 129.71 |
| C10 | CD49a | 0.94 | 3.3e-130 | 121.32 |
| C10 | HLA-DR, DP, DQ | 0.62 | 1.9e-111 | 69.05 |
| C10 | CD29 | 0.43 | 6.2e-136 | 58.01 |
| C10 | HLA-DR | 0.52 | 2.2e-101 | 52.16 |
| C10 | CD319 (CRACC) | 0.37 | 3.8e-97 | 36.04 |
| C10 | CD18 | 0.27 | 3.4e-98 | 26.36 |
| C10 | CD49d | 0.29 | 1.2e-86 | 25.06 |
| C10 | CD279 (PD-1) | 0.40 | 5.7e-59 | 23.28 |
| C10 | CD151 (PETA-3) | 0.34 | 4.3e-57 | 19.10 |
| C10 | CD226 (DNAM-1) | 0.25 | 9.8e-57 | 14.24 |
| C10 | TIGIT (VSTM3) | 0.39 | 2.0e-33 | 12.80 |
| C10 | CD223 (LAG-3) | 0.31 | 1.7e-33 | 10.21 |
| C2 | CD112 (Nectin-2) | 0.50 | 1.4e-23 | 11.48 |

*(continued)*

| Cluster | Marker | log2(fold change) | Adjusted p value | π-score |
|---|---|---|---|---|
| C2 | CD74 | 0.42 | 1.6e-26 | 10.75 |
| C2 | CX3CR1 | 0.44 | 7.8e-22 | 9.19 |
| C2 | CD79b (Igβ) | 0.43 | 1.9e-20 | 8.46 |
| C2 | CD23 | 0.40 | 6.8e-20 | 7.70 |
| C2 | LOX-1 | 0.37 | 1.0e-18 | 6.73 |
| C2 | CD134 (OX40) | 0.39 | 1.6e-16 | 6.23 |
| C2 | CD116 | 0.28 | 3.0e-21 | 5.65 |
| C2 | CD24 | 0.32 | 5.1e-18 | 5.47 |
| C2 | CD303 (BDCA-2) | 0.29 | 3.8e-16 | 4.55 |
| C2 | CD314 (NKG2D) | 0.26 | 8.6e-17 | 4.15 |
| C2 | CD83 | 0.30 | 2.2e-12 | 3.54 |
| C2 | IgE | 0.27 | 2.9e-13 | 3.42 |
| C2 | CD163 | 0.31 | 1.8e-09 | 2.75 |
| C2 | CD141 (Thrombomodulin) | 0.28 | 1.5e-09 | 2.43 |
| C2 | CD275 (B7-H2, B7-RP1, ICOSL) | 0.25 | 3.4e-10 | 2.38 |
| C2 | CD123 | 0.29 | 7.1e-04 | 0.90 |
| C3 | TIGIT (VSTM3) | 0.79 | 3.6e-61 | 47.92 |
| C3 | CD25 | 0.79 | 2.5e-57 | 44.44 |
| C3 | CD95 (Fas) | 0.27 | 2.8e-43 | 11.58 |
| C3 | CD49f | 0.37 | 2.7e-25 | 9.16 |
| C3 | CD194 (CCR4) | 0.32 | 1.2e-20 | 6.34 |
| C4 | CD49b | 0.61 | 5.9e-159 | 96.86 |
| C4 | CD195 (CCR5) | 0.56 | 7.3e-136 | 76.22 |
| C4 | CD161 | 0.69 | 8.3e-108 | 73.66 |
| C4 | CD150 (SLAM) | 0.41 | 1.7e-141 | 57.48 |
| C4 | CD194 (CCR4) | 0.36 | 4.8e-96 | 34.56 |
| C4 | CD49a | 0.42 | 2.8e-81 | 33.52 |
| C4 | CD226 (DNAM-1) | 0.26 | 4.2e-126 | 32.90 |
| C4 | CD25 | 0.33 | 1.2e-96 | 31.57 |
| C4 | CD58 (LFA-3) | 0.26 | 8.3e-119 | 31.27 |
| C4 | CD279 (PD-1) | 0.30 | 8.3e-93 | 27.92 |
| C4 | CD151 (PETA-3) | 0.26 | 4.9e-74 | 19.24 |
| C4 | CD69 | 0.39 | 2.2e-46 | 17.62 |
| C5 | CD49b | 0.56 | 5.3e-182 | 102.14 |
| C5 | CD194 (CCR4) | 0.42 | 1.1e-163 | 67.84 |
| C5 | CD161 | 0.56 | 7.0e-92 | 51.28 |
| C5 | CD29 | 0.27 | 7.6e-187 | 50.77 |
| C5 | CD226 (DNAM-1) | 0.26 | 2.7e-192 | 50.35 |
| C5 | CD58 (LFA-3) | 0.27 | 1.9e-173 | 47.04 |
| C5 | CD150 (SLAM) | 0.32 | 2.3e-145 | 46.34 |
| C5 | CD195 (CCR5) | 0.42 | 1.1e-96 | 40.04 |
| C5 | CD49a | 0.33 | 1.6e-79 | 26.33 |
| C6 | CD55 | 0.44 | 0.0e+00 | Inf |
| C6 | CD45RA | 0.25 | 6.5e-248 | 62.04 |
| C7 | CD38 | 0.33 | 1.8e-177 | 58.69 |
| C7 | CD7 | 0.27 | 4.0e-112 | 30.17 |
| C7 | CD31 | 0.32 | 5.0e-59 | 18.57 |
| C8 | CD38 | 0.38 | 0.0e+00 | Inf |
| C8 | CD7 | 0.35 | 0.0e+00 | Inf |
| C8 | CD27 | 0.27 | 0.0e+00 | Inf |
| C8 | CD31 | 0.45 | 1.9e-203 | 91.10 |
| C8 | CD101 (BB27) | 0.31 | 3.6e-165 | 50.61 |
| C8 | CD183 (CXCR3) | 0.25 | 6.1e-159 | 40.20 |
| C8 | CD9 | 0.29 | 1.0e-50 | 14.26 |

**Supplemental Table 3:** Significant surface markers enriched in HIV+ versus HIV- cells for *in vitro* model.

| Subset | Cluster | Marker | log2(fold change) | Adjusted p value | π-score |
|---|---|---|---|---|---|
| CD4+ | HIV- | CD31 | -1.06 | 1.0e-158 | -167.94 |
| CD4+ | HIV- | CD62L | -0.52 | 9.1e-265 | -136.14 |
| CD4+ | HIV- | CD55 | -0.52 | 1.0e-178 | -91.74 |
| CD4+ | HIV- | CD27 | -0.37 | 1.8e-241 | -89.81 |
| CD4+ | HIV- | CD7 | -0.38 | 5.9e-123 | -46.54 |
| CD4+ | HIV- | CD9 | -0.52 | 8.2e-45 | -23.03 |
| CD4+ | HIV- | CD45RA | -0.22 | 1.9e-89 | -19.14 |
| CD4+ | HIV- | CD38 | -0.27 | 1.4e-44 | -12.02 |
| CD4+ | HIV- | CD3 | -0.06 | 7.5e-83 | -4.54 |
| CD4+ | HIV- | TCR α/β | -0.06 | 8.6e-72 | -4.20 |
| CD4+ | HIV- | CD127 (IL-7Rα) | -0.09 | 4.5e-41 | -3.54 |
| CD4+ | HIV- | CD272 (BTLA) | -0.11 | 1.3e-25 | -2.69 |
| CD4+ | HIV+ | CD195 (CCR5) | 2.08 | 2.2e-308 | 640.57 |
| CD4+ | HIV+ | CD150 (SLAM) | 1.64 | 2.2e-308 | 504.37 |
| CD4+ | HIV+ | CD279 (PD-1) | 1.36 | 1.3e-228 | 309.43 |
| CD4+ | HIV+ | CD49a | 1.68 | 1.7e-172 | 288.33 |
| CD4+ | HIV+ | CD161 | 1.94 | 7.5e-145 | 279.09 |
| CD4+ | HIV+ | CD2 | 0.87 | 2.2e-308 | 267.60 |
| CD4+ | HIV+ | CD95 (Fas) | 0.97 | 5.6e-260 | 251.16 |
| CD4+ | HIV+ | CD49b | 1.61 | 1.6e-154 | 247.25 |
| CD4+ | HIV+ | CD99 | 0.79 | 1.4e-235 | 185.04 |
| CD4+ | HIV+ | CD29 | 1.03 | 1.1e-173 | 177.44 |
| CD4+ | HIV+ | KLRG1 (MAFA) | 1.39 | 4.4e-115 | 158.63 |
| CD4+ | HIV+ | GPR56 | 1.34 | 1.4e-113 | 151.04 |
| CD4+ | HIV+ | HLA-DR, DP, DQ | 1.12 | 2.3e-134 | 149.85 |
| CD4+ | HIV+ | CD151 (PETA-3) | 1.04 | 1.3e-141 | 146.42 |
| CD4+ | HIV+ | CD226 (DNAM-1) | 0.84 | 3.9e-151 | 126.37 |
| CD4+ | HIV+ | CD58 (LFA-3) | 0.89 | 1.4e-121 | 107.82 |
| CD4+ | HIV+ | CD244 (2B4) | 1.21 | 4.7e-89 | 107.21 |
| CD4+ | HIV+ | CD69 | 1.37 | 1.1e-74 | 101.26 |
| CD4+ | HIV+ | HLA-DR | 0.94 | 2.2e-100 | 93.74 |
| CD4+ | HIV+ | CD11a | 0.60 | 3.4e-132 | 79.11 |
| CD4+ | HIV+ | CD196 (CCR6) | 0.99 | 2.2e-66 | 65.06 |
| CD4+ | HIV+ | CD49d | 0.77 | 4.1e-83 | 63.34 |
| CD4+ | HIV+ | CD278 (ICOS) | 0.66 | 1.2e-95 | 62.58 |
| CD4+ | HIV+ | CD45RO | 0.65 | 1.1e-85 | 55.12 |
| CD4+ | HIV+ | CD25 | 0.85 | 1.2e-53 | 45.13 |
| CD4+ | HIV+ | CD192 (CCR2) | 0.91 | 4.4e-49 | 43.98 |
| CD4+ | HIV+ | CD18 | 0.53 | 1.9e-75 | 39.68 |
| CD4+ | HIV+ | CD54 | 0.61 | 2.0e-65 | 39.15 |
| CD4+ | HIV+ | CD26 | 0.70 | 1.9e-53 | 36.72 |
| CD4+ | HIV+ | CD194 (CCR4) | 0.80 | 3.7e-44 | 34.62 |
| CD4+ | HIV+ | CD84 | 0.69 | 3.4e-49 | 33.30 |
| CD4+ | HIV+ | CD146 | 0.84 | 8.1e-40 | 32.93 |
| CD4+ | HIV+ | CD82 | 0.54 | 5.5e-54 | 28.76 |
| CD4+ | HIV+ | CD39 | 0.60 | 9.0e-45 | 26.33 |
| CD4+ | HIV+ | CD119 (IFN-γ R α chain) | 0.60 | 1.7e-35 | 20.78 |
| CD4+ | HIV+ | CD81 (TAPA-1) | 0.52 | 9.3e-24 | 11.99 |
| CD4+ | HIV+ | CD305 (LAIR1) | 0.52 | 4.6e-22 | 11.02 |
| CD4+ | HIV+ | CD352 (NTB-A) | 0.43 | 3.0e-20 | 8.36 |
| CD4+ | HIV+ | CD73 (Ecto-5’-nucleotidase) | 0.63 | 1.3e-13 | 8.05 |
| CD4+ | HIV+ | CD45 | 0.34 | 4.2e-22 | 7.27 |
| CD4+ | HIV+ | CD48 | 0.38 | 4.9e-19 | 7.05 |
| CD4+ | HIV+ | HLA-A,B,C | 0.43 | 2.2e-16 | 6.79 |
| CD4+ | HIV+ | integrin β7 | 0.47 | 3.9e-08 | 3.50 |

*(continued)*

| Subset | Cluster | Marker | log2(fold change) | Adjusted p value | π-score |
|---|---|---|---|---|---|
| CD4+ | HIV+ | CD44 | 0.35 | 2.0e-10 | 3.36 |
| CD4+ | HIV+ | CD183 (CXCR3) | 0.39 | 3.6e-07 | 2.50 |
| CD4+ | HIV+ | CD52 | 0.12 | 5.3e-19 | 2.25 |
| CD4+ | HIV+ | CD43 | 0.30 | 3.3e-07 | 1.94 |
| CD4+ | HIV+ | CD47 | 0.01 | 1.8e-102 | 1.17 |
| CD4+ | HIV+ | CD101 (BB27) | 0.37 | 6.0e-03 | 0.81 |
| CD4+ | HIV+ | CD19 | 0.12 | 2.6e-07 | 0.81 |
| activated/late diff CD4+ | HIV- | CD62L | -0.38 | 1.0e-54 | -20.54 |
| activated/late diff CD4+ | HIV- | CD127 (IL-7Rα) | -0.28 | 1.7e-20 | -5.57 |
| activated/late diff CD4+ | HIV- | CD49f | -0.31 | 4.4e-14 | -4.15 |
| activated/late diff CD4+ | HIV- | CD185 (CXCR5) | -0.26 | 8.6e-13 | -3.09 |
| activated/late diff CD4+ | HIV- | CD194 (CCR4) | -0.18 | 3.1e-08 | -1.33 |
| activated/late diff CD4+ | HIV- | CD27 | -0.06 | 3.7e-14 | -0.77 |
| activated/late diff CD4+ | HIV- | TIGIT (VSTM3) | -0.14 | 7.2e-06 | -0.70 |
| activated/late diff CD4+ | HIV- | CD31 | -0.10 | 2.6e-04 | -0.37 |
| activated/late diff CD4+ | HIV- | CD55 | -0.02 | 3.1e-05 | -0.07 |
| activated/late diff CD4+ | HIV- | CD7 | -0.02 | 8.7e-03 | -0.04 |
| activated/late diff CD4+ | HIV- | CD5 | -0.01 | 1.0e-03 | -0.02 |
| activated/late diff CD4+ | HIV- | CD162 | 0.00 | 1.1e-03 | -0.01 |
| activated/late diff CD4+ | HIV+ | CD195 (CCR5) | 0.90 | 2.1e-32 | 28.34 |
| activated/late diff CD4+ | HIV+ | CD150 (SLAM) | 0.79 | 1.9e-35 | 27.31 |
| activated/late diff CD4+ | HIV+ | CD2 | 0.41 | 4.4e-36 | 14.40 |
| activated/late diff CD4+ | HIV+ | GPR56 | 0.86 | 7.3e-15 | 12.12 |
| activated/late diff CD4+ | HIV+ | CD279 (PD-1) | 0.66 | 1.1e-18 | 11.79 |
| activated/late diff CD4+ | HIV+ | CD95 (Fas) | 0.33 | 3.8e-16 | 5.13 |
| activated/late diff CD4+ | HIV+ | CD119 (IFN-γ R α chain) | 0.46 | 2.5e-10 | 4.46 |
| activated/late diff CD4+ | HIV+ | CD26 | 0.41 | 2.5e-06 | 2.32 |
| activated/late diff CD4+ | HIV+ | HLA-DR, DP, DQ | 0.46 | 2.3e-05 | 2.13 |
| activated/late diff CD4+ | HIV+ | CD244 (2B4) | 0.57 | 2.8e-04 | 2.04 |
| activated/late diff CD4+ | HIV+ | CD11a | 0.27 | 5.9e-08 | 1.97 |
| activated/late diff CD4+ | HIV+ | CD305 (LAIR1) | 0.42 | 2.3e-05 | 1.94 |
| activated/late diff CD4+ | HIV+ | CD146 | 0.55 | 4.1e-04 | 1.88 |
| activated/late diff CD4+ | HIV+ | CD161 | 0.59 | 2.1e-03 | 1.57 |
| activated/late diff CD4+ | HIV+ | CD73 (Ecto-5’-nucleotidase) | 0.56 | 1.9e-03 | 1.53 |
| activated/late diff CD4+ | HIV+ | HLA-DR | 0.43 | 3.4e-04 | 1.49 |
| activated/late diff CD4+ | HIV+ | CD54 | 0.29 | 9.6e-06 | 1.43 |
| activated/late diff CD4+ | HIV+ | CD151 (PETA-3) | 0.38 | 1.6e-03 | 1.07 |
| activated/late diff CD4+ | HIV+ | CD196 (CCR6) | 0.48 | 6.5e-03 | 1.06 |
| activated/late diff CD4+ | HIV+ | CD192 (CCR2) | 0.52 | 1.7e-02 | 0.92 |
| activated/late diff CD4+ | HIV+ | CD48 | 0.23 | 2.5e-04 | 0.82 |
| activated/late diff CD4+ | HIV+ | CD25 | 0.30 | 2.1e-03 | 0.81 |
| activated/late diff CD4+ | HIV+ | CD49b | 0.38 | 9.3e-03 | 0.77 |
| activated/late diff CD4+ | HIV+ | KLRG1 (MAFA) | 0.50 | 3.4e-02 | 0.74 |
| activated/late diff CD4+ | HIV+ | CD49a | 0.43 | 4.5e-02 | 0.58 |
| activated/late diff CD4+ | HIV+ | CD101 (BB27) | 0.39 | 4.1e-02 | 0.53 |
| activated/late diff CD4+ | HIV+ | CD99 | 0.22 | 5.1e-03 | 0.50 |
| activated/late diff CD4+ | HIV+ | CD47 | 0.05 | 5.9e-07 | 0.30 |
| activated/late diff CD4+ | HIV+ | CD52 | 0.07 | 3.0e-03 | 0.17 |
| activated/late diff CD4+ | HIV+ | CD82 | 0.07 | 1.7e-02 | 0.13 |
| early diff CD4+ | HIV- | CD62L | -0.27 | 7.0e-31 | -8.17 |
| early diff CD4+ | HIV- | CD55 | -0.24 | 1.3e-19 | -4.57 |
| early diff CD4+ | HIV- | CD127 (IL-7Rα) | -0.09 | 4.5e-14 | -1.21 |
| early diff CD4+ | HIV- | CD162 | -0.10 | 1.7e-10 | -0.96 |
| early diff CD4+ | HIV- | CD45RA | -0.08 | 1.2e-10 | -0.77 |
| early diff CD4+ | HIV+ | CD278 (ICOS) | 0.57 | 3.0e-17 | 9.46 |
| early diff CD4+ | HIV+ | CD25 | 0.79 | 2.5e-12 | 9.21 |

*(continued)*

| Subset | Cluster | Marker | log2(fold change) | Adjusted p value | π-score |
|---|---|---|---|---|---|
| early diff CD4+ | HIV+ | CD183 (CXCR3) | 0.77 | 8.0e-10 | 7.00 |
| early diff CD4+ | HIV+ | CD11a | 0.55 | 1.1e-12 | 6.53 |
| early diff CD4+ | HIV+ | CD49b | 0.67 | 1.7e-09 | 5.87 |
| early diff CD4+ | HIV+ | CD2 | 0.50 | 3.4e-09 | 4.24 |
| early diff CD4+ | HIV+ | CD38 | 0.60 | 5.4e-07 | 3.74 |
| early diff CD4+ | HIV+ | CD151 (PETA-3) | 0.60 | 1.4e-06 | 3.54 |
| early diff CD4+ | HIV+ | CD305 (LAIR1) | 0.57 | 8.3e-06 | 2.89 |
| early diff CD4+ | HIV+ | CD71 | 0.52 | 2.5e-04 | 1.87 |
| early diff CD4+ | HIV+ | CD18 | 0.45 | 1.1e-04 | 1.80 |
| early diff CD4+ | HIV+ | CD47 | 0.10 | 4.5e-17 | 1.69 |
| early diff CD4+ | HIV+ | CD101 (BB27) | 0.57 | 1.6e-03 | 1.61 |
| early diff CD4+ | HIV+ | HLA-A,B,C | 0.18 | 5.3e-07 | 1.10 |
| early diff CD4+ | HIV+ | CD95 (Fas) | 0.42 | 5.6e-03 | 0.95 |
| early diff CD4+ | HIV+ | CD4 | 0.42 | 1.1e-02 | 0.82 |
| early diff CD4+ | HIV+ | CD19 | 0.12 | 3.9e-02 | 0.16 |
| early diff CD4+ | HIV+ | CD49f | 0.04 | 3.0e-03 | 0.10 |

**Supplemental Table 4:** Differential peak list for activated CD4+ cells between HIV+ and HIV- for *in vitro* model.

| Subset | Chr | Start | End | Log2FC | FDR | π-score | In Gene | Nearest TSS | Nearest TSS Dist |
|---|---|---|---|---|---|---|---|---|---|
| HIV- | chrX | 133763071 | 133763571 | -5.76 | 2.0e-02 | -9.73 | GPC3 | GPC3 | 222324 |
| HIV- | chr11 | 128840083 | 128840583 | -2.32 | 3.4e-04 | -8.04 | KCNJ1 | KCNJ1 | 26790 |
| HIV- | chr3 | 120344302 | 120344802 | -1.62 | 4.3e-04 | -5.47 | LRRC58 | LRRC58 | 4537 |
| HIV- | chr16 | 29013010 | 29013510 | -1.51 | 4.3e-04 | -5.08 | MIR3680-1 | LAT | 28184 |
| HIV- | chr5 | 75078009 | 75078509 | -2.44 | 9.5e-03 | -4.94 | ANKRD31 | LINC01336 | -25166 |
| HIV- | chr17 | 62164986 | 62165486 | -3.33 | 4.4e-02 | -4.51 | - | MED13 | -99704 |
| HIV- | chr13 | 73984895 | 73985395 | -1.91 | 7.8e-03 | -4.03 | KLF12 | KLF12 | 148510 |
| HIV- | chr7 | 127322273 | 127322773 | -1.49 | 2.1e-03 | -4.00 | - | GRM8 | -68979 |
| HIV- | chr6 | 149366167 | 149366667 | -1.44 | 1.8e-03 | -3.93 | TAB2 | SUMO4 | -33560 |
| HIV- | chr1 | 87045694 | 87046194 | -2.48 | 2.7e-02 | -3.89 | HS2ST1, LINC01140 | LINC01140 | 52685 |
| HIV- | chr5 | 109921610 | 109922110 | -1.41 | 2.0e-03 | -3.82 | - | LINC01848 | 38428 |
| HIV- | chr5 | 39186867 | 39187367 | -1.84 | 8.4e-03 | -3.82 | FYB1 | FYB1 | 87161 |
| HIV- | chr10 | 3800532 | 3801032 | -2.74 | 4.7e-02 | -3.65 | - | KLF6 | -15251 |
| HIV- | chr1 | 39653445 | 39653945 | -1.90 | 1.7e-02 | -3.35 | - | HEYL | -13500 |
| HIV- | chr3 | 10263496 | 10263996 | -2.19 | 3.0e-02 | -3.34 | TATDN2 | TATDN2 | 15473 |
| HIV- | chr8 | 120472266 | 120472766 | -2.41 | 4.5e-02 | -3.25 | MTBP | MRPL13 | -26864 |
| HIV- | chr7 | 1925895 | 1926395 | -1.53 | 7.8e-03 | -3.23 | MAD1L1 | MIR4655 | -81642 |
| HIV- | chrX | 136674455 | 136674955 | -1.51 | 7.8e-03 | -3.18 | ARHGEF6 | CD40LG | 26262 |
| HIV- | chr12 | 64494944 | 64495444 | -1.89 | 2.1e-02 | -3.17 | TBK1 | TBK1 | 43064 |
| HIV- | chr15 | 44560876 | 44561376 | -1.87 | 2.1e-02 | -3.15 | EIF3J | EIF3J | 23819 |
| HIV- | chr8 | 120775213 | 120775713 | -2.19 | 3.8e-02 | -3.13 | SNTB1, LOC101927543 | LOC101927543 | 13960 |
| HIV- | chr11 | 14463556 | 14464056 | -2.33 | 4.7e-02 | -3.10 | COPB1 | COPB1 | 35971 |
| HIV- | chr6 | 37289865 | 37290365 | -2.29 | 4.6e-02 | -3.06 | TBC1D22B | TMEM217 | -31710 |
| HIV- | chr15 | 88904581 | 88905081 | -1.01 | 2.0e-03 | -2.72 | MFGE8 | MFGE8 | 8330 |
| HIV- | chr3 | 153643435 | 153643935 | -1.59 | 2.1e-02 | -2.68 | - | C3orf79 | 158940 |
| HIV- | chr16 | 29827058 | 29827558 | -1.85 | 4.4e-02 | -2.52 | BOLA2, SLX1A, SLX1A-SULT1A3, SULT1A4, LOC388242, PAGR1, MVP | MVP | 6664 |
| HIV- | chr2 | 203987082 | 203987582 | -1.65 | 4.4e-02 | -2.24 | - | ICOS | 50334 |
| HIV- | chr3 | 187027350 | 187027850 | -1.03 | 7.8e-03 | -2.18 | ST6GAL1 | ST6GAL1 | 96865 |
| HIV- | chr10 | 118732697 | 118733197 | -1.59 | 4.8e-02 | -2.09 | CACUL1 | CACUL1 | 22052 |
| HIV- | chr3 | 118955786 | 118956286 | -1.29 | 4.1e-02 | -1.78 | IGSF11 | IGSF11-AS1 | 12713 |
| HIV- | chr13 | 49371836 | 49372336 | -1.12 | 3.1e-02 | -1.70 | CAB39L | CAB39L | 71790 |
| HIV- | chr14 | 32102444 | 32102944 | -1.21 | 4.5e-02 | -1.63 | ARHGAP5 | ARHGAP5-AS1 | -25651 |
| HIV- | chr10 | 43332102 | 43332602 | -1.16 | 4.0e-02 | -1.62 | - | FXYD4 | -39040 |
| HIV- | chr1 | 100927241 | 100927741 | -0.91 | 3.0e-02 | -1.38 | SLC30A7 | SLC30A7 | 31165 |

*(continued)*

| Subset | Chr | Start | End | Log2FC | FDR | π-score | In Gene | Nearest TSS | Nearest TSS Dist |
|---|---|---|---|---|---|---|---|---|---|
| HIV- | chr17 | 66331088 | 66331588 | -0.94 | 3.4e-02 | -1.38 | PRKCA | PRKCA | 28452 |
| HIV- | chr14 | 89903512 | 89904012 | -1.01 | 4.7e-02 | -1.34 | FOXN3, EFCAB11 | FOXN3 | 50647 |
| HIV- | chr13 | 51825061 | 51825561 | -0.85 | 2.7e-02 | -1.33 | TMEM272 | TMEM272 | 19614 |
| HIV- | chr16 | 27235313 | 27235813 | -0.99 | 4.6e-02 | -1.32 | MIR3680-1, NSMCE1 | KDM8 | 31818 |
| HIV- | chr10 | 110225646 | 110226146 | -0.83 | 2.6e-02 | -1.32 | MXI1 | MXI1 | 18041 |
| HIV- | chr13 | 40485590 | 40486090 | -0.61 | 1.5e-02 | -1.10 | LINC00598 | LINC00598 | 49717 |
| HIV+ | chr1 | 221829595 | 221830095 | 1.39 | 3.9e-05 | 6.15 | LINC01655 | LINC01655 | 10571 |
| HIV+ | chr9 | 73869213 | 73869713 | 2.09 | 1.4e-03 | 5.99 | - | LOC101927358 | -501818 |
| HIV+ | chr6 | 117465645 | 117466145 | 2.53 | 6.7e-03 | 5.50 | GOPC, DCBLD1 | DCBLD1 | 11828 |
| HIV+ | chr14 | 68719698 | 68720198 | 1.16 | 3.5e-05 | 5.17 | - | ZFP36L1 | 76055 |
| HIV+ | chr8 | 72000175 | 72000675 | 1.52 | 1.4e-03 | 4.34 | MSC-AS1 | TRPA1 | 74942 |
| HIV+ | chr14 | 59275611 | 59276111 | 1.76 | 5.1e-03 | 4.05 | DAAM1 | DAAM1 | 86965 |
| HIV+ | chr15 | 101199696 | 101200196 | 1.15 | 3.3e-04 | 4.00 | CHSY1 | CHSY1 | 51736 |
| HIV+ | chr9 | 75148484 | 75148984 | 2.03 | 1.1e-02 | 3.99 | - | OSTF1 | 59941 |
| HIV+ | chr15 | 90412628 | 90413128 | 2.94 | 4.4e-02 | 3.98 | IQGAP1 | IQGAP1 | 24410 |
| HIV+ | chr6 | 13747556 | 13748056 | 1.10 | 4.3e-04 | 3.70 | - | RANBP9 | -35992 |
| HIV+ | chr2 | 101223824 | 101224324 | 1.52 | 5.4e-03 | 3.44 | TBC1D8 | CNOT11 | -28478 |
| HIV+ | chr2 | 136362680 | 136363180 | 1.86 | 1.5e-02 | 3.38 | - | CXCR4 | -244515 |
| HIV+ | chr3 | 71447689 | 71448189 | 2.51 | 4.6e-02 | 3.36 | FOXP1 | MIR1284 | 93900 |
| HIV+ | chr9 | 86348418 | 86348918 | 1.66 | 9.7e-03 | 3.35 | TUT7 | TUT7 | 5536 |
| HIV+ | chr7 | 23595009 | 23595509 | 2.50 | 4.7e-02 | 3.32 | - | CCDC126 | -1870 |
| HIV+ | chr11 | 14288180 | 14288680 | 1.57 | 9.1e-03 | 3.22 | RRAS2 | RRAS2 | 75826 |
| HIV+ | chr3 | 71392986 | 71393486 | 1.90 | 2.0e-02 | 3.22 | FOXP1 | FOXP1-AS1 | 103217 |
| HIV+ | chr17 | 3795234 | 3795734 | 1.80 | 2.0e-02 | 3.04 | ITGAE | ITGAE | 5509 |
| HIV+ | chr1 | 25099439 | 25099939 | 2.11 | 3.8e-02 | 3.00 | - | MIR4425 | 75936 |
| HIV+ | chr11 | 43316418 | 43316918 | 2.09 | 3.8e-02 | 2.97 | API5 | API5 | 4455 |
| HIV+ | chr13 | 27131122 | 27131622 | 0.85 | 3.8e-04 | 2.89 | USP12 | USP12-AS1 | -31233 |
| HIV+ | chr5 | 72194799 | 72195299 | 2.07 | 4.0e-02 | 2.88 | MAP1B | MIR4803 | 25332 |
| HIV+ | chr4 | 26076133 | 26076633 | 1.52 | 1.5e-02 | 2.75 | - | RBPJ | -86822 |
| HIV+ | chr16 | 82660155 | 82660655 | 2.05 | 4.8e-02 | 2.70 | CDH13 | MIR8058 | -28276 |
| HIV+ | chr15 | 38610792 | 38611292 | 1.30 | 9.1e-03 | 2.66 | - | RASGRP1 | -45217 |
| HIV+ | chr8 | 133152859 | 133153359 | 1.20 | 7.0e-03 | 2.58 | - | WISP1 | -37680 |
| HIV+ | chr8 | 28365700 | 28366200 | 1.93 | 4.8e-02 | 2.53 | ZNF395 | PNOC | 48714 |
| HIV+ | chr10 | 22505996 | 22506496 | 1.85 | 4.5e-02 | 2.48 | - | SPAG6 | 160551 |
| HIV+ | chr11 | 34235166 | 34235666 | 1.43 | 2.0e-02 | 2.41 | ABTB2 | ABTB2 | 122342 |
| HIV+ | chr4 | 121256332 | 121256832 | 1.18 | 9.5e-03 | 2.39 | - | TNIP3 | -28866 |
| HIV+ | chr16 | 23504680 | 23505180 | 1.09 | 6.7e-03 | 2.37 | MIR3680-1, GGA2 | GGA2 | 16815 |
| HIV+ | chr5 | 127785555 | 127786055 | 1.07 | 8.4e-03 | 2.22 | CCDC192 | CCDC192 | 82165 |
| HIV+ | chr8 | 97980061 | 97980561 | 1.27 | 2.0e-02 | 2.14 | MATN2 | SNORA72 | 61656 |
| HIV+ | chr10 | 22597125 | 22597625 | 1.60 | 4.7e-02 | 2.13 | PIP4K2A | PIP4K2A | 116930 |

*(continued)*

| Subset | Chr | Start | End | Log2FC | FDR | π-score | In Gene | Nearest TSS | Nearest TSS Dist |
|---|---|---|---|---|---|---|---|---|---|
| HIV+ | chr2 | 65232542 | 65233042 | 1.24 | 2.0e-02 | 2.10 | ACTR2 | ACTR2 | 4789 |
| HIV+ | chr11 | 128551675 | 128552175 | 1.22 | 2.2e-02 | 2.03 | ETS1 | ETS1-AS1 | 25533 |
| HIV+ | chr13 | 50245013 | 50245513 | 1.42 | 3.9e-02 | 2.01 | DLEU1 | DLEU2 | -119293 |
| HIV+ | chr3 | 46369194 | 46369694 | 1.43 | 4.1e-02 | 1.97 | LOC102724297 | CCR5 | -1160 |
| HIV+ | chr10 | 88270827 | 88271327 | 1.03 | 1.2e-02 | 1.96 | - | RNLS | 313203 |
| HIV+ | chr6 | 108921393 | 108921893 | 1.19 | 2.4e-02 | 1.94 | ARMC2 | ARMC2-AS1 | 2210 |
| HIV+ | chr1 | 32263726 | 32264226 | 1.42 | 4.5e-02 | 1.91 | LCK | LCK | 12487 |
| HIV+ | chr8 | 100493218 | 100493718 | 1.33 | 4.7e-02 | 1.76 | - | ANKRD46 | 66066 |
| HIV+ | chr4 | 101039328 | 101039828 | 1.24 | 4.1e-02 | 1.73 | PPP3CA | EMCN | -159202 |
| HIV+ | chr2 | 178514591 | 178515091 | 0.78 | 6.4e-03 | 1.71 | PLEKHA3 | TTN-AS1 | -6092 |
| HIV+ | chr3 | 155862523 | 155863023 | 1.29 | 4.8e-02 | 1.70 | - | GMPS | -7513 |
| HIV+ | chr14 | 74573941 | 74574441 | 1.08 | 4.3e-02 | 1.48 | LTBP2 | LTBP2 | 37937 |
| HIV+ | chr22 | 39961397 | 39961897 | 0.87 | 2.0e-02 | 1.47 | GRAP2 | FAM83F | -33052 |
| HIV+ | chr13 | 76991309 | 76991809 | 0.65 | 1.3e-02 | 1.21 | CLN5 | CLN5 | 649 |
| HIV+ | chr8 | 66467771 | 66468271 | 0.76 | 4.1e-02 | 1.06 | ADHFE1, VXN | VXN | 7768 |

**Supplemental Table 5:** Significant surface markers enriched in each cluster (over all other cells) during treated infection.

| Cluster | Marker | log2(fold change) | Adjusted p value | π-score |
|---|---|---|---|---|
| C1 | KLRG1 (MAFA) | 0.28 | 5.4e-263 | 72.25 |
| C10 | CD74 | 0.50 | 0.0e+00 | Inf |
| C10 | CD79b (Igβ) | 0.41 | 0.0e+00 | Inf |
| C10 | CD200 (OX2) | 0.35 | 0.0e+00 | Inf |
| C10 | CD23 | 0.35 | 0.0e+00 | Inf |
| C10 | IgE | 0.25 | 0.0e+00 | Inf |
| C11 | CD27 | 0.50 | 0.0e+00 | Inf |
| C11 | CD7 | 0.47 | 0.0e+00 | Inf |
| C11 | integrin β7 | 0.33 | 0.0e+00 | Inf |
| C11 | CD161 | 0.28 | 0.0e+00 | Inf |
| C12 | IgM | 0.57 | 0.0e+00 | Inf |
| C12 | Ig light chain λ | 0.48 | 0.0e+00 | Inf |
| C12 | CD39 | 0.43 | 0.0e+00 | Inf |
| C12 | CD62L | 0.41 | 0.0e+00 | Inf |
| C12 | Ig light chain κ | 0.31 | 0.0e+00 | Inf |
| C12 | CD49b | 0.29 | 0.0e+00 | Inf |
| C12 | CD57 | 0.28 | 0.0e+00 | Inf |
| C12 | CD55 | 0.25 | 0.0e+00 | Inf |
| C13 | CD74 | 0.53 | 0.0e+00 | Inf |
| C13 | CD79b (Igβ) | 0.42 | 0.0e+00 | Inf |
| C13 | CD23 | 0.37 | 0.0e+00 | Inf |
| C13 | CD200 (OX2) | 0.35 | 0.0e+00 | Inf |
| C13 | CD55 | 0.27 | 0.0e+00 | Inf |
| C13 | IgE | 0.25 | 0.0e+00 | Inf |
| C14 | CD161 | 0.49 | 0.0e+00 | Inf |
| C14 | CD26 | 0.49 | 0.0e+00 | Inf |
| C14 | CD192 (CCR2) | 0.41 | 0.0e+00 | Inf |
| C14 | CD55 | 0.41 | 0.0e+00 | Inf |
| C14 | CD49f | 0.38 | 0.0e+00 | Inf |
| C14 | CD73 (Ecto-5'-nucleotidase) | 0.37 | 0.0e+00 | Inf |
| C14 | CD7 | 0.33 | 0.0e+00 | Inf |
| C14 | CD27 | 0.33 | 0.0e+00 | Inf |
| C14 | CD127 (IL-7Rα) | 0.32 | 0.0e+00 | Inf |
| C14 | CD69 | 0.32 | 0.0e+00 | Inf |
| C14 | CD150 (SLAM) | 0.29 | 0.0e+00 | Inf |
| C14 | CD49d | 0.27 | 0.0e+00 | Inf |
| C14 | CD35 | 0.26 | 7.2e-60 | 15.42 |
| C15 | CD49b | 0.35 | 0.0e+00 | Inf |
| C15 | CD39 | 0.29 | 0.0e+00 | Inf |
| C15 | CD9 | 0.28 | 0.0e+00 | Inf |
| C15 | HLA-DR | 0.25 | 0.0e+00 | Inf |
| C16 | CD69 | 0.51 | 0.0e+00 | Inf |
| C16 | CD194 (CCR4) | 0.35 | 0.0e+00 | Inf |
| C16 | CD7 | 0.32 | 0.0e+00 | Inf |
| C16 | CD49f | 0.29 | 0.0e+00 | Inf |
| C16 | CD5 | 0.27 | 0.0e+00 | Inf |
| C16 | CD272 (BTLA) | 0.26 | 0.0e+00 | Inf |
| C16 | CD49a | 0.32 | 2.6e-260 | 84.29 |
| C18 | CD74 | 0.82 | 0.0e+00 | Inf |
| C18 | CD79b (Igβ) | 0.80 | 0.0e+00 | Inf |
| C18 | IgE | 0.79 | 0.0e+00 | Inf |
| C18 | CD23 | 0.74 | 0.0e+00 | Inf |
| C18 | LOX-1 | 0.67 | 0.0e+00 | Inf |
| C18 | CD134 (OX40) | 0.67 | 0.0e+00 | Inf |
| C18 | CD314 (NKG2D) | 0.67 | 0.0e+00 | Inf |

*(continued)*

| Cluster | Marker | log2(fold change) | Adjusted p value | π-score |
|---|---|---|---|---|
| C18 | CD200 (OX2) | 0.65 | 0.0e+00 | Inf |
| C18 | CD24 | 0.64 | 0.0e+00 | Inf |
| C18 | CD83 | 0.62 | 0.0e+00 | Inf |
| C18 | CD163 | 0.60 | 0.0e+00 | Inf |
| C18 | CD45RA | 0.59 | 0.0e+00 | Inf |
| C18 | CX3CR1 | 0.58 | 0.0e+00 | Inf |
| C18 | CD93 | 0.55 | 0.0e+00 | Inf |
| C18 | CD141 (Thrombomodulin) | 0.53 | 0.0e+00 | Inf |
| C18 | CD303 (BDCA-2) | 0.52 | 0.0e+00 | Inf |
| C18 | CD116 | 0.50 | 0.0e+00 | Inf |
| C18 | CD112 (Nectin-2) | 0.50 | 0.0e+00 | Inf |
| C18 | CD275 (B7-H2, B7-RP1, ICOSL) | 0.49 | 0.0e+00 | Inf |
| C18 | CD13 | 0.46 | 0.0e+00 | Inf |
| C18 | CD223 (LAG-3) | 0.46 | 0.0e+00 | Inf |
| C18 | CD122 (IL-2Rβ) | 0.44 | 0.0e+00 | Inf |
| C18 | CD115 (CSF-1R) | 0.44 | 0.0e+00 | Inf |
| C18 | Podoplanin | 0.43 | 0.0e+00 | Inf |
| C18 | FcεRIα | 0.43 | 0.0e+00 | Inf |
| C18 | CD183 (CXCR3) | 0.40 | 0.0e+00 | Inf |
| C18 | CD142 | 0.40 | 0.0e+00 | Inf |
| C18 | CD196 (CCR6) | 0.39 | 0.0e+00 | Inf |
| C18 | CD85j (ILT2) | 0.39 | 0.0e+00 | Inf |
| C18 | NKp80 | 0.38 | 0.0e+00 | Inf |
| C18 | CD158 (KIR2DL1/S1/S3/S5) | 0.38 | 0.0e+00 | Inf |
| C18 | CD109 | 0.37 | 0.0e+00 | Inf |
| C18 | CD14 | 0.37 | 0.0e+00 | Inf |
| C18 | CD169 (Sialoadhesin, Siglec-1) | 0.36 | 0.0e+00 | Inf |
| C18 | CD185 (CXCR5) | 0.32 | 0.0e+00 | Inf |
| C18 | CD152 (CTLA-4) | 0.32 | 0.0e+00 | Inf |
| C18 | TIGIT (VSTM3) | 0.31 | 0.0e+00 | Inf |
| C18 | CD354 (TREM-1) | 0.31 | 0.0e+00 | Inf |
| C18 | CD137 (4-1BB) | 0.29 | 0.0e+00 | Inf |
| C18 | CD94 | 0.29 | 0.0e+00 | Inf |
| C18 | CD195 (CCR5) | 0.28 | 0.0e+00 | Inf |
| C18 | CD158e1 (KIR3DL1, NKB1) | 0.28 | 0.0e+00 | Inf |
| C18 | CD38 | 0.28 | 0.0e+00 | Inf |
| C18 | CD279 (PD-1) | 0.28 | 0.0e+00 | Inf |
| C18 | Ig light chain λ | 0.25 | 0.0e+00 | Inf |
| C19 | CD26 | 0.26 | 1.2e-181 | 46.66 |
| C19 | CD192 (CCR2) | 0.25 | 3.6e-140 | 34.86 |
| C2 | CD244 (2B4) | 0.41 | 2.8e-291 | 118.05 |
| C2 | GPR56 | 0.44 | 9.4e-239 | 104.72 |
| C2 | CD195 (CCR5) | 0.31 | 4.7e-219 | 68.20 |
| C2 | KLRG1 (MAFA) | 0.34 | 1.9e-203 | 68.10 |
| C2 | CD57 | 0.32 | 1.3e-104 | 33.67 |
| C20 | CD11c | 1.63 | 0.0e+00 | Inf |
| C20 | CD33 | 1.36 | 0.0e+00 | Inf |
| C20 | CD31 | 1.24 | 0.0e+00 | Inf |
| C20 | CLEC12A | 1.22 | 0.0e+00 | Inf |
| C20 | HLA-DR | 1.17 | 0.0e+00 | Inf |
| C20 | HLA-DR, DP, DQ | 0.98 | 0.0e+00 | Inf |
| C20 | CD116 | 0.91 | 0.0e+00 | Inf |
| C20 | CD141 (Thrombomodulin) | 0.87 | 0.0e+00 | Inf |
| C20 | CD93 | 0.86 | 0.0e+00 | Inf |
| C20 | CD41 | 0.85 | 0.0e+00 | Inf |
| C20 | CD61 | 0.79 | 0.0e+00 | Inf |

*(continued)*

| Cluster | Marker | log2(fold change) | Adjusted p value | π-score |
|---|---|---|---|---|
| C20 | CD9 | 0.77 | 0.0e+00 | Inf |
| C20 | CD63 | 0.76 | 0.0e+00 | Inf |
| C20 | CD328 (Siglec-7) | 0.74 | 0.0e+00 | Inf |
| C20 | CD32 | 0.69 | 0.0e+00 | Inf |
| C20 | CD88 (C5aR) | 0.62 | 0.0e+00 | Inf |
| C20 | CD85j (ILT2) | 0.53 | 0.0e+00 | Inf |
| C20 | CD35 | 0.86 | 3.6e-266 | 227.61 |
| C20 | CLEC1B (CLEC2) | 0.68 | 1.7e-295 | 199.86 |
| C20 | CD11b | 0.56 | 6.2e-295 | 163.63 |
| C20 | CD39 | 0.60 | 8.3e-240 | 142.56 |
| C20 | CD49b | 0.57 | 2.8e-182 | 102.94 |
| C20 | CD354 (TREM-1) | 0.43 | 4.5e-230 | 97.59 |
| C20 | CD62P (P-Selectin) | 0.47 | 4.1e-206 | 95.73 |
| C20 | CD14 | 0.53 | 1.3e-170 | 89.92 |
| C20 | CD38 | 0.49 | 5.1e-171 | 83.43 |
| C20 | CD244 (2B4) | 0.42 | 7.2e-191 | 79.93 |
| C20 | CD13 | 0.36 | 6.6e-190 | 68.39 |
| C20 | CD151 (PETA-3) | 0.39 | 1.1e-166 | 64.33 |
| C20 | CD101 (BB27) | 0.40 | 2.0e-142 | 56.77 |
| C20 | CD119 (IFN-γ R α chain) | 0.32 | 2.0e-136 | 43.55 |
| C20 | CD54 | 0.34 | 1.5e-83 | 28.49 |
| C20 | CD305 (LAIR1) | 0.25 | 1.3e-48 | 11.98 |
| C20 | CD42b | 0.27 | 1.2e-34 | 9.21 |
| C20 | FcεRIα | 0.37 | 1.3e-22 | 8.16 |
| C3 | CD56 (NCAM) | 1.21 | 0.0e+00 | Inf |
| C3 | CD94 | 1.21 | 0.0e+00 | Inf |
| C3 | CD314 (NKG2D) | 0.89 | 0.0e+00 | Inf |
| C3 | CD244 (2B4) | 0.89 | 0.0e+00 | Inf |
| C3 | CD16 | 0.87 | 0.0e+00 | Inf |
| C3 | CD11c | 0.83 | 0.0e+00 | Inf |
| C3 | GPR56 | 0.80 | 0.0e+00 | Inf |
| C3 | NKp80 | 0.62 | 0.0e+00 | Inf |
| C3 | CD63 | 0.60 | 0.0e+00 | Inf |
| C3 | CD9 | 0.59 | 0.0e+00 | Inf |
| C3 | CD57 | 0.80 | 1.3e-222 | 176.97 |
| C3 | CD31 | 0.58 | 3.2e-248 | 144.62 |
| C3 | CD61 | 0.57 | 2.5e-252 | 143.88 |
| C3 | CD8 | 0.63 | 1.5e-217 | 137.58 |
| C3 | CD41 | 0.60 | 9.9e-229 | 135.78 |
| C3 | CLEC1B (CLEC2) | 0.47 | 5.7e-168 | 78.86 |
| C3 | CD122 (IL-2Rβ) | 0.38 | 6.0e-199 | 75.54 |
| C3 | CD151 (PETA-3) | 0.38 | 2.1e-198 | 75.03 |
| C3 | CD328 (Siglec-7) | 0.50 | 1.5e-144 | 71.75 |
| C3 | CD11b | 0.30 | 1.5e-233 | 70.71 |
| C3 | CD32 | 0.31 | 2.7e-222 | 68.41 |
| C3 | TIGIT (VSTM3) | 0.39 | 5.1e-152 | 58.31 |
| C3 | CD38 | 0.41 | 2.1e-126 | 51.95 |
| C3 | HLA-DR | 0.31 | 3.5e-122 | 37.23 |
| C3 | CD49b | 0.33 | 4.4e-38 | 12.14 |
| C4 | GPR56 | 0.92 | 0.0e+00 | Inf |
| C4 | CD57 | 0.83 | 0.0e+00 | Inf |
| C4 | KLRG1 (MAFA) | 0.74 | 0.0e+00 | Inf |
| C4 | CX3CR1 | 0.51 | 0.0e+00 | Inf |
| C4 | CD74 | 0.43 | 0.0e+00 | Inf |
| C4 | CD244 (2B4) | 0.38 | 0.0e+00 | Inf |
| C4 | CD122 (IL-2Rβ) | 0.35 | 0.0e+00 | Inf |

*(continued)*

| Cluster | Marker | log2(fold change) | Adjusted p value | π-score |
|---|---|---|---|---|
| C4 | CD79b (Igβ) | 0.34 | 0.0e+00 | Inf |
| C4 | IgM | 0.28 | 0.0e+00 | Inf |
| C4 | CD23 | 0.26 | 0.0e+00 | Inf |
| C4 | CD305 (LAIR1) | 0.26 | 0.0e+00 | Inf |
| C5 | GPR56 | 0.76 | 0.0e+00 | Inf |
| C5 | KLRG1 (MAFA) | 0.72 | 0.0e+00 | Inf |
| C5 | CD244 (2B4) | 0.51 | 0.0e+00 | Inf |
| C5 | CD195 (CCR5) | 0.43 | 0.0e+00 | Inf |
| C5 | CD57 | 0.42 | 0.0e+00 | Inf |
| C5 | HLA-DR, DP, DQ | 0.41 | 0.0e+00 | Inf |
| C5 | CD162 | 0.34 | 0.0e+00 | Inf |
| C5 | HLA-DR | 0.31 | 0.0e+00 | Inf |
| C5 | CD49d | 0.28 | 0.0e+00 | Inf |
| C5 | CD151 (PETA-3) | 0.28 | 0.0e+00 | Inf |
| C5 | CD11a | 0.26 | 0.0e+00 | Inf |
| C6 | CD57 | 0.90 | 0.0e+00 | Inf |
| C6 | KLRG1 (MAFA) | 0.79 | 0.0e+00 | Inf |
| C6 | GPR56 | 0.76 | 0.0e+00 | Inf |
| C6 | CX3CR1 | 0.54 | 0.0e+00 | Inf |
| C6 | CD39 | 0.36 | 0.0e+00 | Inf |
| C6 | CD122 (IL-2Rβ) | 0.31 | 0.0e+00 | Inf |
| C6 | Ig light chain λ | 0.27 | 0.0e+00 | Inf |
| C6 | CD244 (2B4) | 0.27 | 0.0e+00 | Inf |
| C6 | IgM | 0.26 | 0.0e+00 | Inf |
| C6 | CD85j (ILT2) | 0.26 | 0.0e+00 | Inf |
| C7 | CD83 | 0.59 | 0.0e+00 | Inf |
| C7 | CD303 (BDCA-2) | 0.56 | 0.0e+00 | Inf |
| C7 | CD223 (LAG-3) | 0.55 | 0.0e+00 | Inf |
| C7 | CD24 | 0.52 | 0.0e+00 | Inf |
| C7 | CD275 (B7-H2, B7-RP1, ICOSL) | 0.51 | 0.0e+00 | Inf |
| C7 | NKp80 | 0.48 | 0.0e+00 | Inf |
| C7 | IgE | 0.47 | 0.0e+00 | Inf |
| C7 | CD169 (Sialoadhesin, Siglec-1) | 0.47 | 0.0e+00 | Inf |
| C7 | CD134 (OX40) | 0.46 | 0.0e+00 | Inf |
| C7 | CD23 | 0.45 | 0.0e+00 | Inf |
| C7 | CD116 | 0.45 | 0.0e+00 | Inf |
| C7 | CD137 (4-1BB) | 0.43 | 0.0e+00 | Inf |
| C7 | Podoplanin | 0.43 | 0.0e+00 | Inf |
| C7 | CD142 | 0.42 | 0.0e+00 | Inf |
| C7 | LOX-1 | 0.42 | 0.0e+00 | Inf |
| C7 | CD112 (Nectin-2) | 0.42 | 0.0e+00 | Inf |
| C7 | CD109 | 0.41 | 0.0e+00 | Inf |
| C7 | CD115 (CSF-1R) | 0.38 | 0.0e+00 | Inf |
| C7 | CD85j (ILT2) | 0.38 | 0.0e+00 | Inf |
| C7 | CD163 | 0.38 | 0.0e+00 | Inf |
| C7 | CD152 (CTLA-4) | 0.35 | 0.0e+00 | Inf |
| C7 | CD79b (Igβ) | 0.34 | 0.0e+00 | Inf |
| C7 | CD74 | 0.33 | 0.0e+00 | Inf |
| C7 | Ig light chain κ | 0.33 | 0.0e+00 | Inf |
| C7 | CD200 (OX2) | 0.32 | 0.0e+00 | Inf |
| C7 | CX3CR1 | 0.32 | 0.0e+00 | Inf |
| C7 | CD304 (Neuropilin-1) | 0.32 | 0.0e+00 | Inf |
| C7 | CD314 (NKG2D) | 0.31 | 0.0e+00 | Inf |
| C7 | FcεRIα | 0.31 | 0.0e+00 | Inf |
| C7 | TIGIT (VSTM3) | 0.30 | 0.0e+00 | Inf |
| C7 | CD45RA | 0.29 | 0.0e+00 | Inf |

*(continued)*

| Cluster | Marker | log2(fold change) | Adjusted p value | π-score |
|---|---|---|---|---|
| C7 | CD141 (Thrombomodulin) | 0.29 | 0.0e+00 | Inf |
| C7 | CD158e1 (KIR3DL1, NKB1) | 0.27 | 0.0e+00 | Inf |
| C7 | CD14 | 0.27 | 0.0e+00 | Inf |
| C7 | Ig light chain λ | 0.26 | 0.0e+00 | Inf |
| C7 | CD267 (TACI) | 0.26 | 0.0e+00 | Inf |
| C7 | CD93 | 0.26 | 0.0e+00 | Inf |
| C7 | CD37 | 0.25 | 0.0e+00 | Inf |
| C9 | CD25 | 0.70 | 0.0e+00 | Inf |
| C9 | CD39 | 0.66 | 0.0e+00 | Inf |
| C9 | TIGIT (VSTM3) | 0.57 | 0.0e+00 | Inf |
| C9 | CD71 | 0.53 | 0.0e+00 | Inf |
| C9 | HLA-DR, DP, DQ | 0.51 | 0.0e+00 | Inf |
| C9 | HLA-DR | 0.47 | 0.0e+00 | Inf |
| C9 | CD194 (CCR4) | 0.39 | 0.0e+00 | Inf |
| C9 | CD27 | 0.29 | 0.0e+00 | Inf |
| C9 | CD278 (ICOS) | 0.27 | 0.0e+00 | Inf |
| C9 | CD101 (BB27) | 0.39 | 1.2e-264 | 102.94 |

**Supplemental Table 6:** Significant surface markers enriched in HIV+ versus HIV- cells during treated infection.

| Subset | Cluster | Marker | log2(fold change) | Adjusted p value | π-score |
|---|---|---|---|---|---|
| CD4+ (DESeq2) | HIV- | CD200 (OX2) | -0.84 | 1.4e-09 | -7.42 |
| CD4+ (DESeq2) | HIV- | CD74 | -0.75 | 2.0e-09 | -6.49 |
| CD4+ (DESeq2) | HIV- | CD41 | -0.80 | 6.2e-07 | -4.98 |
| CD4+ (DESeq2) | HIV- | CD314 (NKG2D) | -0.64 | 5.5e-07 | -4.02 |
| CD4+ (DESeq2) | HIV- | GPR56 | -0.87 | 4.0e-04 | -2.96 |
| CD4+ (DESeq2) | HIV- | CD158e1 (KIR3DL1, NKB1) | -0.69 | 5.4e-05 | -2.94 |
| CD4+ (DESeq2) | HIV- | CD103 (Integrin αE) | -0.71 | 1.9e-04 | -2.65 |
| CD4+ (DESeq2) | HIV- | CD3 | -0.18 | 6.5e-13 | -2.19 |
| CD4+ (DESeq2) | HIV- | CD8 | -0.55 | 1.7e-03 | -1.54 |
| CD4+ (DESeq2) | HIV- | CD62L | -0.29 | 1.7e-05 | -1.40 |
| CD4+ (DESeq2) | HIV- | CD79b (Igβ) | -0.55 | 3.2e-03 | -1.38 |
| CD4+ (DESeq2) | HIV- | CD122 (IL-2Rβ) | -0.34 | 9.9e-04 | -1.01 |
| CD4+ (DESeq2) | HIV- | CD49b | -0.45 | 1.3e-02 | -0.84 |
| CD4+ (DESeq2) | HIV- | CD305 (LAIR1) | -0.32 | 2.5e-03 | -0.83 |
| CD4+ (DESeq2) | HIV- | CD38 | -0.51 | 2.3e-02 | -0.83 |
| CD4+ (DESeq2) | HIV- | IgE | -0.40 | 1.1e-02 | -0.79 |
| CD4+ (DESeq2) | HIV- | CD94 | -0.43 | 2.1e-02 | -0.72 |
| CD4+ (DESeq2) | HIV- | LOX-1 | -0.33 | 1.6e-02 | -0.59 |
| CD4+ (DESeq2) | HIV- | CD134 (OX40) | -0.31 | 1.9e-02 | -0.53 |
| CD4+ (DESeq2) | HIV- | CD56 (NCAM) | -0.37 | 4.3e-02 | -0.51 |
| CD4+ (DESeq2) | HIV- | TCR α/β | -0.03 | 1.1e-02 | -0.07 |
| CD4+ (DESeq2) | HIV+ | CD99 | 0.36 | 7.6e-05 | 1.49 |
| CD4+ (DESeq2) | HIV+ | CD2 | 0.34 | 8.4e-03 | 0.70 |
| CD4+ (Wilcoxon) | HIV- | CD200 (OX2) | -0.18 | 2.0e-03 | -0.49 |
| CD4+ (Wilcoxon) | HIV- | CD74 | -0.18 | 3.7e-02 | -0.26 |
| CD4+ (Wilcoxon) | HIV+ | CD150 (SLAM) | 0.18 | 6.1e-07 | 1.10 |
| CD4+ (Wilcoxon) | HIV+ | CD192 (CCR2) | 0.18 | 1.7e-04 | 0.70 |
| CD4+ (Wilcoxon) | HIV+ | CD49d | 0.13 | 4.4e-04 | 0.44 |
| CD4+ (Wilcoxon) | HIV+ | CD26 | 0.14 | 5.6e-03 | 0.32 |
| CD4+ (Wilcoxon) | HIV+ | HLA-DR, DP, DQ | 0.16 | 2.2e-02 | 0.27 |
| CD4+ (Wilcoxon) | HIV+ | CD279 (PD-1) | 0.14 | 1.9e-02 | 0.24 |
| CD4+ (Wilcoxon) | HIV+ | HLA-DR | 0.13 | 3.3e-02 | 0.20 |
| Tcm/Ttm (DESeq2) | HIV- | CD62L | -0.53 | 9.9e-03 | -1.06 |
| Tcm/Ttm (DESeq2) | HIV+ | CD71 | 1.10 | 6.7e-04 | 3.48 |
| Tcm/Ttm (DESeq2) | HIV+ | CD2 | 0.46 | 2.4e-02 | 0.75 |
| Tcm/Ttm (DESeq2) | HIV+ | CD11a | 0.47 | 3.8e-02 | 0.68 |
| Tem (DESeq2) | HIV- | CD74 | -1.25 | 2.1e-05 | -5.87 |
| Tem (DESeq2) | HIV- | CD41 | -1.29 | 9.9e-04 | -3.88 |
| Tem (DESeq2) | HIV- | CD3 | -0.39 | 3.2e-09 | -3.29 |
| Tem (DESeq2) | HIV- | CD158e1 (KIR3DL1, NKB1) | -1.41 | 5.5e-03 | -3.19 |
| Tem (DESeq2) | HIV- | CD71 | -1.05 | 9.9e-04 | -3.15 |
| Tem (DESeq2) | HIV- | CD101 (BB27) | -1.11 | 1.9e-03 | -3.01 |
| Tem (DESeq2) | HIV- | GPR56 | -0.88 | 2.2e-03 | -2.32 |
| Tem (DESeq2) | HIV- | CD9 | -1.02 | 1.1e-02 | -2.01 |
| Tem (DESeq2) | HIV- | CD61 | -0.99 | 1.7e-02 | -1.75 |
| Tem (DESeq2) | HIV- | CD314 (NKG2D) | -0.93 | 1.5e-02 | -1.69 |
| Tem (DESeq2) | HIV- | CD305 (LAIR1) | -0.66 | 2.7e-03 | -1.68 |
| Tem (DESeq2) | HIV- | CD56 (NCAM) | -0.95 | 3.2e-02 | -1.42 |
| Tem (DESeq2) | HIV- | CD200 (OX2) | -0.96 | 3.4e-02 | -1.41 |
| Tem (DESeq2) | HIV- | CD27 | -0.61 | 1.9e-02 | -1.06 |
| Tem (DESeq2) | HIV- | CD151 (PETA-3) | -0.45 | 4.6e-02 | -0.60 |
| Tem (DESeq2) | HIV- | CD226 (DNAM-1) | -0.20 | 4.7e-02 | -0.27 |
| Tem (DESeq2) | HIV+ | CD49a | 1.58 | 1.7e-03 | 4.38 |
| Tem (DESeq2) | HIV+ | CD195 (CCR5) | 0.93 | 4.2e-02 | 1.28 |

**Supplemental Table 7:** Autologous virus sequences used in study.

| Individual | Fasta Prefix or Name | Location | Publication |
|---|---|---|---|
| A08 | chrA08 | GenBank; Supplementary File 1 | Gondim et al., 2021 |
| A09 | chrA09 | GenBank; Supplementary File 1 | Gondim et al., 2021 |
| A01 | chrA01. | Supplementary File 1 | This paper |
| B45 | chrBEAT2045_ | Supplementary File 1 | This paper |
| ND492 | SUMA_TF1 | GenBank (JN944928.1) | Ochsenbauer et al., 2012 |
